# The translation initiation factor DHX29 appears to pull on mRNA in a direction opposite to scanning

**DOI:** 10.1101/2025.07.13.664561

**Authors:** Da Cui, Tatyana Pestova, Christopher Hellen, Amedee des Georges

## Abstract

The DExH-box helicase DHX29 plays a critical role in mammalian translation initiation. It is required for the scanning of mRNAs with complex 5’UTRs. Despite its importance, the detailed mechanism of DHX29’s action has remained debated. Using structural models derived from AlphaFold and cryo-EM structure of DHX29 bound to the ribosomal 43S pre-initiation complex, we provide a revised structural framework that clarifies the interplay between DHX29, the 40S ribosomal subunit, and eIF3. Our findings suggest that the 40S subunit regulates DHX29’s NTPase activity through an activation mechanism resembling the G-patch protein regulation of DEAH helicases. Moreover, our model supports a 3’ to 5’ translocase mechanism, in which DHX29 transiently pulls the mRNA opposite to the scanning direction, destabilizing stable stem-loops trapped in the mRNA channel and allowing scanning to resume. This structural analysis refines our understanding of DHX29’s function and provides new hypotheses regarding its role in mRNA unwinding during scanning and start codon selection.

## Introduction

The regulation of protein synthesis is key to cellular homeostasis. In eukaryotes, initiation is a rate-limiting step of translation, whose regulation allows for a rapid and localized response to stimuli. For most mRNAs, translation initiation occurs via a scanning mechanism whereby a scanning-competent 43S preinitiation complex (43S PIC), composed of the 40S subunit and eukaryotic initiation factors (eIFs) 1, 1A, 3, and 2 bound to initiator Met-tRNA_i_^Met^, attaches to the capped 5’ end of an mRNA before scanning for the AUG start codon (Jackson, 2010). Scanning involves unwinding the secondary structures of the mRNA 5’ untranslated region (5’-UTR) by eIF4A, 4B, and 4F. While the eIF4F complex was shown to impose 3’-directionality during scanning (Skabkin, 2013), there is evidence of retrograde 3’-5’ scanning (Abaeva, 2016; Gu, 2021; Li, 2022; Wang, 2022), shown by in vitro reconstitution to occur in the absence of these factors and facilitated by eIF1 and eIF1A (Abaeva, 2016). eIF1 and eIF1A have been shown to open the mRNA channel to facilitate scanning and play an important role in ensuring the fidelity of start codon recognition (Pestova, 1998; Battiste, 2000; Maag, 2006; Passmore, 2007; Nanda, 2009; Saini, 2010).

In mammals, a large portion of mRNAs contains stable stem-loops in the 5’-UTR that play important roles in the regulation of gene expression. Scanning through such structured mRNAs requires the additional factor DHX29, a DEAH box RNA helicase of the RNA helicase A (RHA) subfamily (Pisareva, 2008). Its silencing disrupts general translation, impairing the formation of polysomes and leading to the accumulation of mRNA-free 80S ribosomes, most likely due to the ribosome’s inability to scan through those structured mRNAs (Parsyan, 2009).

DHX29 comprises a helicase core whose architecture is conserved throughout DExH helicases and an N-terminal region (NTR) unique to DHX29 (Pisareva, 2008). The helicase core is formed by recombinase A (RecA)-like domains 1 and 2, and a C-terminal region (CTR) composed of a winged-helix (WH) domain, a ratchet-like domain, and an oligonucleotide binding (OB) domain. Cryo-electron microscopy (cryo-EM) structures show that DHX29 binds near the mRNA entry channel of the 40S subunit (Hashem, 2013; des Georges, 2015). The helicase core interacts primarily with the tip of helix 16 (h16) of the 18S rRNA, bridging h16 with the beak, as well as with the b and i subunits of eIF3 (Dhote, 2012; Hashem 2013; des Georges 2015). The unique 534aa-long N-terminal region (NTR), which bridges from the helicase core positioned at the mRNA entrance to the intersubunit face of the 40S subunit, includes a conserved dsRNA-binding domain (dsRBD) that binds to the ribosome next to the A-site and is linked via a long alpha helix to a ubiquitin-associated-like (UBA-like) domain. The UBA-like domain connects the NTR to the helicase core by interacting with the CTR (Dhote, 2012; Hashem, 2013; des Georges 2015; Sweeney, 2021).

DExH helicases bind and hydrolyze NTPs via conserved motifs in their RecA1 and RecA2 domains (Pyle, 2008). Motifs I, II, V, and VI are involved in NTP binding and hydrolysis (Walker, 1982; Smith and Rayment, 1996; Leipe, 2003; Tauchert, 2017). Motifs Ia, Ib, and IV are involved in RNA/DNA binding and translocation (Gorbalenya and Koonin, 1993; Weng, 1996; Fernández, 1997; Korolev, 1998), and motif III has been implicated in coupling NTPase and unwinding activities (Tauchert, 2017).

While DHX29 has not been shown to have helicase activity on its own, NTPase activity is necessary for its function in promoting 48S formation on mRNAs with stable stems in the 5’UTR (Pisareva, 2008; Dhote, 2012). The NTPase activity of DHX29 is enhanced more than 10-fold by the binding of the 40S subunit (Pisareva, 2008) by an unknown mechanism. U_70_ or (CUUU)_9_ RNA also enhances DHX29 NTPase activity to a lesser extent. NTPase activity is highest in the presence of the 40S subunit and (CUUU)_9_ RNA (Pisareva, 2008).

While distant from the helicase core, deletion of dsRBD in the NTR suppresses the NTPase stimulation by the 40S subunit. This influence on the NTPase activity appears to be geometric given that the lengthening of the long helix that connects this domain to the helicase core is critical to that function (Sweeney,2021). This strongly suggests that DHX29 promotes the unwinding of the mRNA stem-loops by acting on the conformation of the ribosome, but direct unwinding by its helicase domain cannot be entirely discounted. The NTPase activity of DHX29 is also affected by a large insert between motifs V and VI of the RecA2 domain whose sequence is specific to DHX29 (**Figure 1A**). It was shown to perform an autoinhibitory role on the intrinsic NTPase activity of the helicase domain. This autoinhibition is relieved when DHX29 binds to the ribosome or U_70_ RNA (Dhote, 2012). The mechanism by which the ribosome and the NTR stimulate DHX29 NTPase activity and how the large insert inhibits it is still not understood at the molecular level.

**Figure 1:**
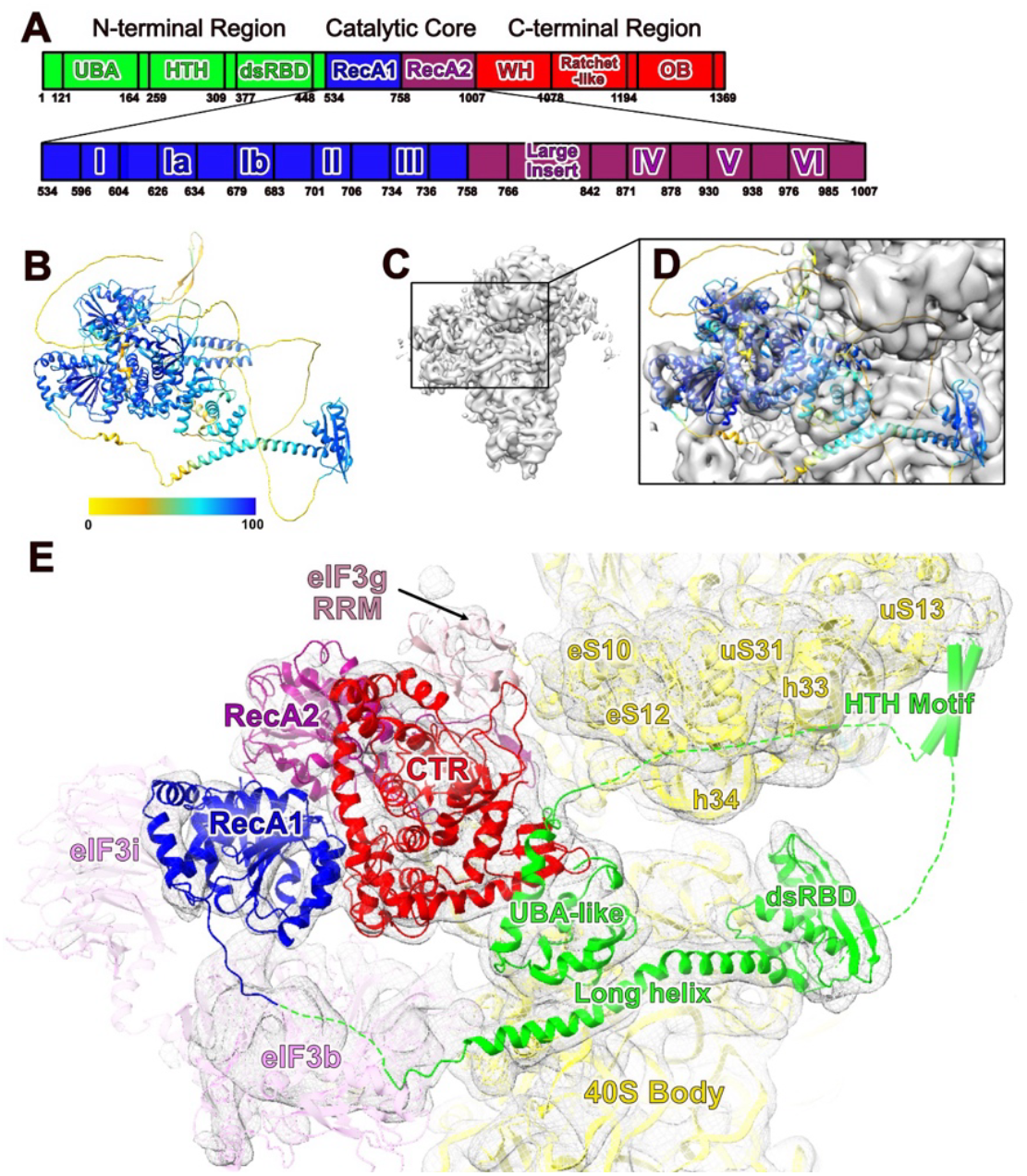
Model of human DHX29 in the context of 43S PIC derived from Alphafold2. **A)** Domain organization of human DHX29 **B**) The atomic model of human DHX29 predicted by Alphafold2 colored by pLDDT as a measure of confidence level **C**) Cryo-EM density of DHX29-bound mammalian 43S Pre-initiation complex (EMD-3057). **D**) Rigid body fit of the predicted model of human DHX29 into the cryo-EM density prior to adjustments. **E**) Refined model of human DHX29.

Knowledge derived from other DExH family helicases, such as Prp43 and Prp22, may provide insights into how DHX29’s NTPase activity is transduced into direct or indirect helicase activity. When mRNA is not associated with the DExH helicases Prp22, Prp43 and DHX9, the mRNA channel has been observed in an open conformation and the position of the RecA2 domain is dynamic (He, 2010; He, 2017; Tauchert, 2017; Hamann, 2019; Lee, 2023). For Prp22, and Prp43, the binding of mRNA into the mRNA channel enhances NTPase activity by stabilizing the RecA2 domain into a catalytically active position where RecA1 and RecA2 domains are in close contact to form the NTP binding pocket and catalytic site (He, 2010; He, 2017; Tauchert, 2017; Hamann, 2019). G-patch proteins, found to be activators of several DExH box RNA helicases, can further stabilize this catalytically active conformation, including inducing conformational change of the RNA tunnel (Becker, 2023) and constriction of movement of the RecA domains which in turn enhances the NTPase activity of DExH-box RNA helicases (Studer, 2020). During translocation, conformational changes of the CTR regulate the opening and closing of the mRNA channel, which is formed by the RecA, ratchet, and OB domains (He, 2010; Walbott, 2010). A hook-turn motif in the RecA1 domain is key to the 3’-5’ translocation process (Tauchert, 2017; Hamann 2019). A key feature of the DEAH helicase Prp43 is the large opening of the mRNA binding channel by the rearrangement of the C-terminal domain, which appears key to the binding of complex folded RNA (Tauchert, 2017). While such a feature may be conserved in other DExH box helicases, we do not know whether it is also conserved in DHX29, or whether similar large rearrangements of the C-terminal domain of DHX29 could be coupled to ribosome conformational changes.

Recently, AlphaFold2 (Jumper, 2021) predicted the structure of human DHX29 with a high confidence level for most parts. This model agrees strikingly well with the cryo-EM structure of DHX29 bound to the 43S PIC (des Georges 2015; Sweeney 2021). Here we refined this model based on the cryo-EM map of the DHX29-bound 43S (EMD 3057), which allowed us to shed new light on the architecture and interactions of DHX29 with 43S PIC and the regulation of its NTPase activity. Our model shows that residues important for the translocase activity of DEAH family helicases are conserved, pointing to a direct role in unwinding mRNA stem loops. This leads us to hypothesize that DHX29 uses a similar winching mechanism as its paralogs to unwind mRNA stem-loops, taking footing on the ribosome to pull on the mRNA in an opposite direction to scanning and destabilize stem-loops. In addition, h16 of the ribosome interacts with the helicase core via an interface that is equivalent to that of the DExH stimulating proteins of the G-patch family, pointing to a similar NTPase activation mechanism. We also observe the large insert slotted into the mRNA entrance channel, which may play a role in positioning the RecA2 domain in a productive conformation. Finally, we observe that the NTR of DHX29 forms multiple interactions with the ribosome beak, implying that both direct helicase activity and control of the ribosome conformational dynamics are important for DHX29’s function in scanning highly structured mRNAs.

## Materials and Methods

### MODEL BUILDING

Alphafold2 (Jumper, 2021) was used to predict the structure of human DHX29 (Uniprot ID: Q7Z478). The regions with low pLDDT scores were removed, and the rest were separated into the domains of NTR, RecA1, RecA2, and CTT, which were then fitted individually into the previously determined cryo-EM structure (EMD-3057; des Georges, 2015) and then joined. The combined model was refined in Phenix real space refinement with the original AF2 model as the reference-based restraint. The atomic model of rabbit late-stage 48S complex(PDB: 6YAM) was separated into different modules: the 40S head together with the cap-binding motif of eIF3d, the 40S body, the eIF2•GTP/Met-tRNA_i_^Met^ ternary complex, and the octamer core of eIF3, which were fitted into the pre-determined cryo-EM density (EMD-3057) individually by rigid-body fitting in chimeraX (Goddard, 2017), and then manually combined in Coot (Emsley, 2004). The atomic models of eIF3e and eIF3d NTT were replaced by the models predicted by Alphafold3 (Abramson, 2024) and were fitted into the cryo-EM map by rigid body with the N-terminal residues manually removed. The model of human eIF3g RRM (PDB: 6MZW) was fitted into the density next to DHX29 by rigid body fitting with the constraint that its position should be able to maintain its interaction with eS10. For eiF3 peripheral domains eIF3a CTT, eIF3b and eIF3i, the model predicted by Alphafold3 was used due to its better fit to the map than the initial rabbit eIF3 peripheral domains structure (PDB: 5A5U).

The C-terminal residues of uS19, uS13, and uS9 were removed as corresponding densities are not visible in the cryo-EM map. Alphafold2 prediction of residues 207-226 of uS2 was added to the atomic model of uS2 without further refinement. The model of eS26 CTT was replaced by that of the crystal structure of the rabbit 40S subunit (PDB: 4KZY) without any modification.

The models of the missing 18S ribosomal RNA residue 241-269 (helix 9), residue 753-783 (eS6S), and residue 1756-1764 (helix 44) were predicted by Alphafold3 (Abramson, 2024), fitted to EMD-3057 by rigid body, and real-space refined in Phenix (Afonine, 2018).

### MODEL VALIDATION

The model statistics, the cross-correlation between the model and the map, and the map-to-model FSC, are calculated at Phenix (Terwilliger, 2022). The EMRinger score was calculated in the EMRinger module in Phenix (Barad, 2015; Terwilliger, 2022). Given the map resolution of 9Å, all sidechains were removed from the deposited model.

### ELECTROSTATIC POTENTIAL CALCULATION

The electrostatic potential discussed in this study is calculated via the Adaptive Poisson-Boltzmann Solver (Jurrus, 2018) implemented in Pymol.

### SEQUENCE ALIGNMENTS

Protein sequences of DHX29 from 21 species, spanning from yeast to *Chordata*, were retrieved from Uniparc (Leinonen, 2004). Multiple sequence alignment was performed using Clustal Omega (RRID: SCR_001591) (Sievers, 2017), employing default parameters. The resulting alignments were visualized using Jalview (Waterhouse, 2009). To assess sequence conservation, an entropy-based measure was calculated using AL2CO (Pei, 2001), and the conservation scores were mapped onto the DHX29 structure using ChimeraX (Goddard, 2017).

### STRUCTURAL COMPARISONS

Structures of different DExH-box RNA helicases were aligned to the RecA1 domain using Matchmaker (Meng, 2006), which is implemented in chimeraX.

## Results

AlphaFold2 generated a model for DHX29 with a predicted local distance difference test (pLDDT) above 80% for most structured regions of the protein (Uniprot ID: Q7Z478; **Figure 1B**), which represents a high per-residue confidence in residue positions (Jumper, 2021). Most of the structured domains of the DHX29 model predicted by AlphaFold2 (**Figure 1B**) fitted readily into the cryo-EM map of the DHX29-bound 43S PIC (EMD-3057, **Figure 1C-D**). Only the regions with pLDDT values below 50, which is considered a good predictor of intrinsic disorder (Mariani, 2013), were not readily accounted for in the density. Minor rearrangements of domain orientations improved the fitting of all domains of DHX29 (**Figure S1A-S1D**), improving the cross-correlation to 0.74 (**Table 1**). The EMRinger score of 0.2 reflects the expected lack of side chain rotamer detail for a map in the 6–10 Å resolution range. Accordingly, our model analysis is limited to Cα backbone positions. Overall, the DHX29-bound 43S map provides experimental evidence for domain and secondary structure positions from residues 82 to 173, 371 to 517, 541 to 778, and 827 to 1369, representing around 74% of the residues of the human DHX29 (**Table 1**). In addition, we do observe densities corresponding to a linker that extends from the UBA-like domain (which we call linker-I), the first two helices of a putative HTH motif, a linker that extends between the HTH motif and the dsRBD (which we call linker-II), and the complete large insert in RecA2 domain (**Figure 1E**). The remaining sequence appears to be unstructured.

**Table 1:**
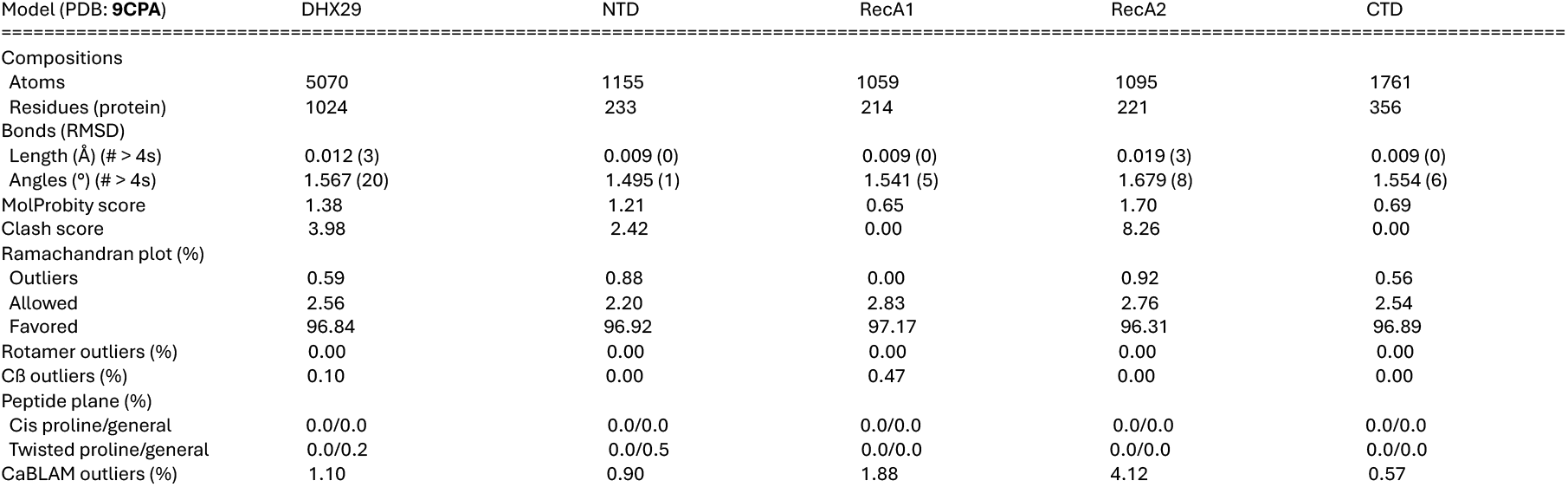
Model statistics of human DHX29 and its domains. The statistics are generated in Phenix against the map EMD-3057.

### THE NTR FORMS MULTIPLE INTERACTIONS WITH THE RIBOSOME BEAK

In the NTR of the protein, in addition to the UBA-like domain, the putative dsRNA binding domain (dsRBD), and the long helix connecting the dsRBD and RecA1 domain (Sweeney, 2021), we could observe a low-resolution density on the 40S head next to uS13 and uS19 and adjacent to initiator tRNA and the A-site (**Figure 2A**). That density shows evidence for two alpha-helices of the helix-turn-helix (HTH) motif predicted by AlphaFold (**Figure 2B, D**). We observe at a lower resolution an elongated density connecting it to the dsRBD domain (**Figure 1, 2A**), which further supports the assignment of those low-resolution densities. Still, the resolution in this part of the map is too low to precisely and unambiguously assign the position and orientation of the HTH motif. A third helix with a lower pLDDT score is also present in the AlphaFold model but not predicted to interact with the other two and we cannot assign density for it. It is interesting to note that HTH motifs generally act as recognition motifs for double-stranded DNA or RNA, which hints that this motif may serve to recruit DHX29 when RNA hairpins are present in the A-site (Brennan, 1989).

**Figure 2:**
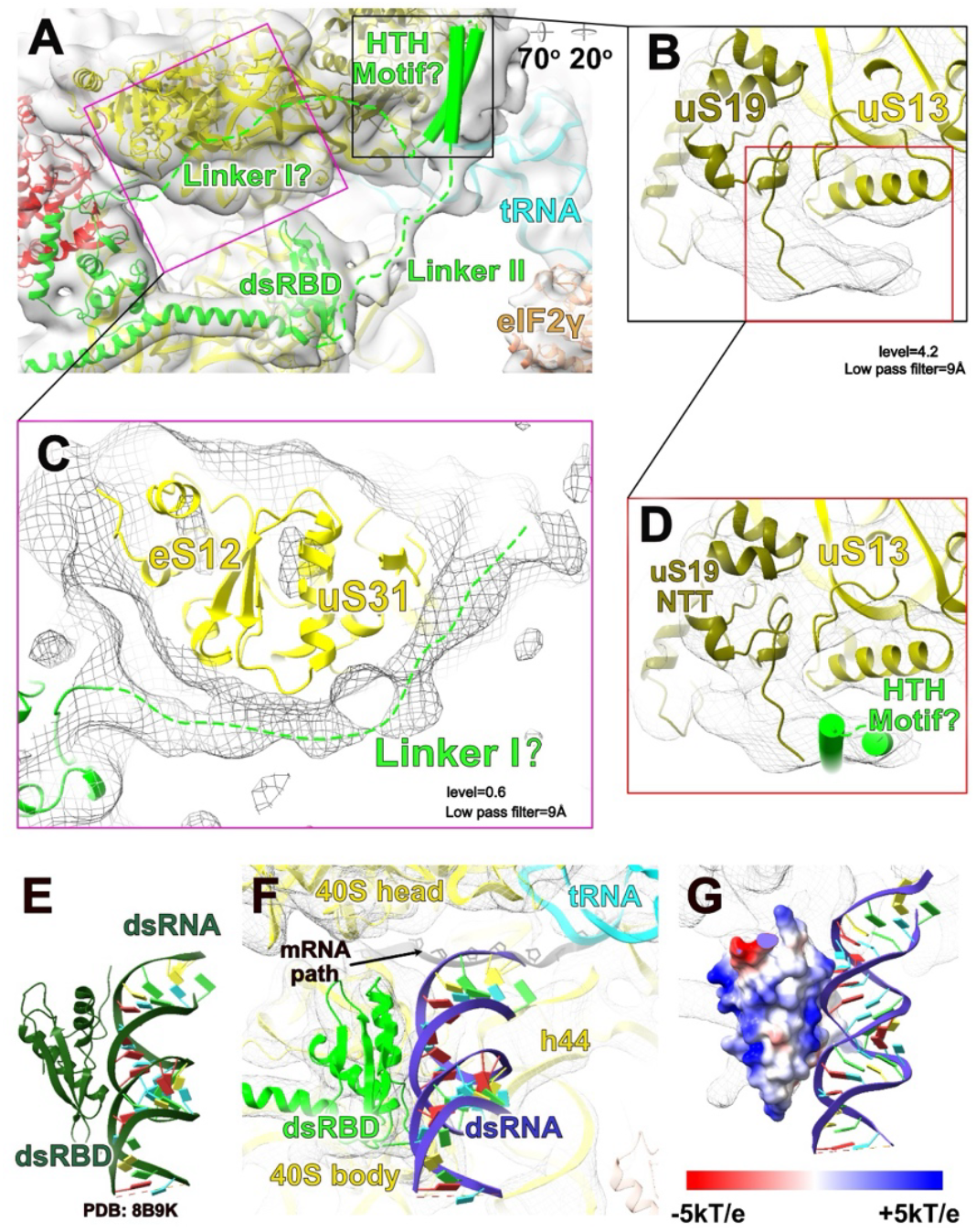
Tentative orientation determination of the ribosomal interaction with the HTH and flexible linker in the NTR. **A**) Topology of the NTR of human DHX29 **B**) A top view of the density of the HTH motif **C**) The linker extends from the UBA-like domain in the NTR and interacts with the ribosomal proteins eS12 and uS31. **D**) A top view of the HTH motif showing the helical shape of the density with the HTH model fitting in. **E**) The atomic model of dsRBD interaction with dsRNA of Drosophila melanogaster MLE **F)** Docking of dsRNA model (PDB: 8B9K) into the model of human DHX29. **G)** Electrostatic surface representation of the dsRBD bound to dsRNA. A positively charged region on the dsRBD (blue) is observed at the putative interface with dsRNA.

The density connecting the HTH motif and the dsRBD in our structure indicates they are linked by a flexible region, suggesting a functional connection between the two. A recent cryo-EM structure of MLE showed that its N-terminal dsRBD directly interacts with dsRNA, confirming its role in dsRNA recognition (Jatgap, 2023). Given the similarities in structure and shape between the dsRBDs of MLE and human DHX29 (**Figure 2E, 2F**), we propose that DHX29’s dsRBD similarly engages with dsRNA, specifically the mRNA stem-loops that can occupy the A site. This idea is supported by the positively charged electrostatic surface on the dsRBD of DHX29 (**Figure 2G**) and the lack of steric clashes with components of the 43S PIC (**Figure 2F**).

Furthermore, since stem-loops can stall mRNA scanning when blocked in the A site (Abaeva et al., 2011), we hypothesize that both the dsRBD and the HTH motif recognize and bind these stalled stem-loops just upstream of the decoding center. Consequently, these interactions would likely assist in recruiting DHX29 to the ribosomal mRNA entry channel to facilitate the unwinding of stem-loops. We can also observe several low-resolution densities connecting the N-terminal helix and UBA-like domain with the beak, including an elongated density running from the tip of the beak towards the HTH motif (**Figure 2C and S2**). The unstructured polypeptides extending from the N-terminal helix and the C-terminal end of the UBA-like domain are likely candidates for these densities. The sequence of these polypeptides is conserved (∼90% conservation for residues 54-86 and ∼50% conservation for residues 167-193; Sweeney, 2021) and deleting residues 54 to 86 in the N-terminal of the first helix abrogates DHX29’s activity (Sweeney, 2021). It is tempting to speculate that these bridges between the UBA-like domain and the ribosome beak play an important functional role, either in constraining the topology of the mRNA entry channel or the conformation of the ribosome head or even possibly in transducing force generated by the helicase core NTPase activity to influence ribosome head conformation. More work will be needed to test these hypotheses.

### EIF3G APPEARS TO FORM PART OF THE DHX29 MRNA ENTRANCE CHANNEL

The cooperation between eIF3 and DHX29 has been suggested to be important for the unwinding of mRNA stem-loops during scanning (Pisareva, 2016), which is partially supported by the interactions between the RecA1 domain and the two β-propellers of eIF3b and eIF3i previously described (des Georges 2015). We also observe a low-resolution density between the RecA2 domain and the ribosome head in the DHX29-bound 43S map (EMD: 3057; des Georges, 2015). We tentatively assign this density to the eIF3g RNA-recognition motif (RRM) (**Figure 3C and S3**), based on its size and the similar location it has in the human 48S and in the 43S open state, where it was observed to interact with h16, uS3, and eS10 CTT in the absence of DHX29 (Brito Querido, 2020; Kratzat, 2021). In the DHX29-bound 43S PIC, we observe that the eIF3g RRM still interacts with the 40S subunit via the CTT of eS10 but no longer with h16 and uS3 (**Figure 3B, C, and S3**), suggesting a change in position of eIF3g upon DHX29 association.

**Figure 3:**
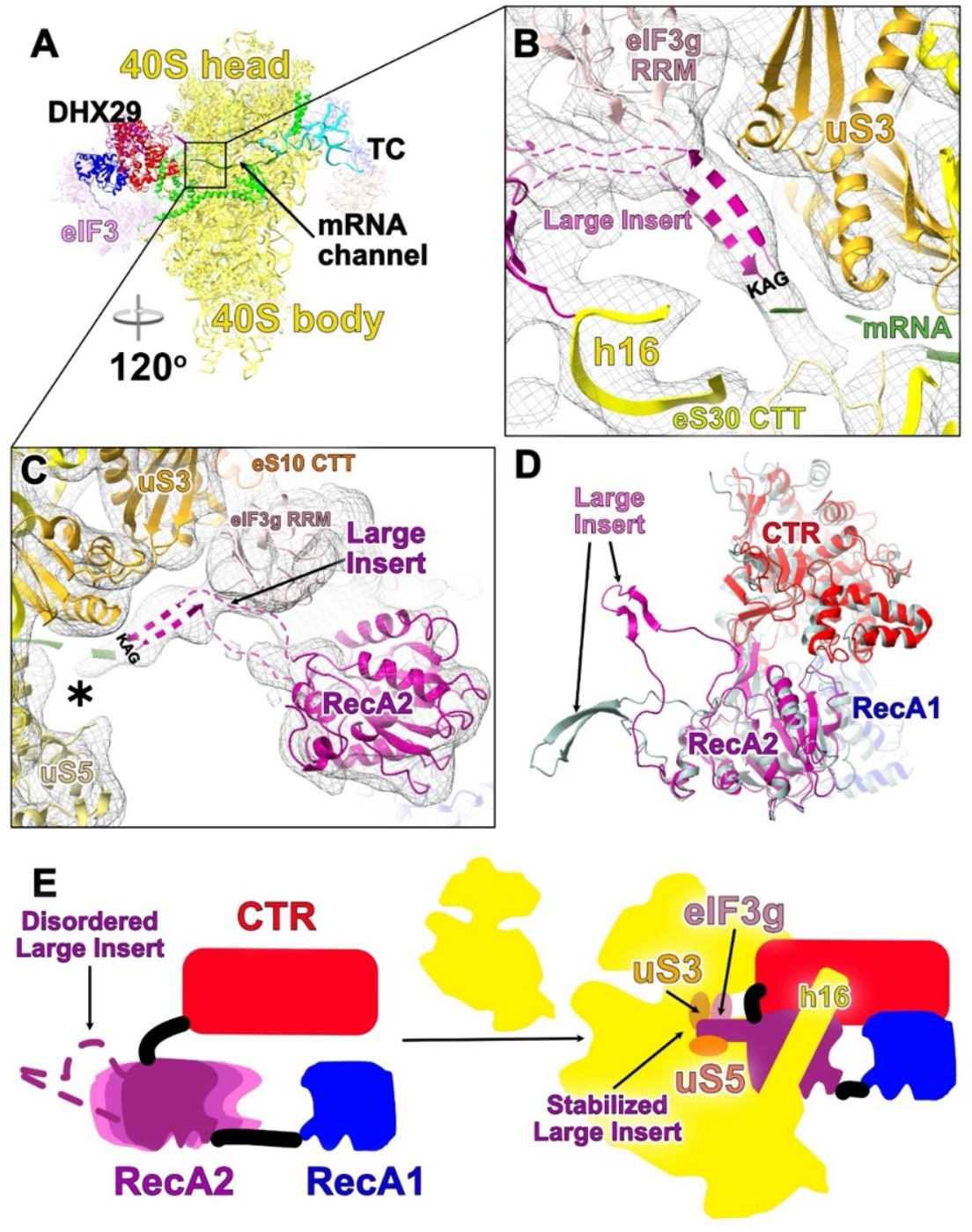
The large insert is positioned at the 40S ribosome mRNA entrance. **A**) Overview of DHX29-bound mammalian 43S PIC. The helicase core of DHX29 is located at the ribosomal mRNA entrance channel while the NTR extends to the vicinity of the decoding center. **B**) Proposed position of the large insert β-hairpin loop at the entrance of the mRNA channel, its conserved KAG residues close to the mRNA path. Dashed ribbons depict the hypotetical nature of the position. The mRNA path is represented as **green dashes** according to its position in the 48S complex (PDB: 6ZMW). **C**) Large insert position relative to eIF3g RRM and uS3 in the ribosomal mRNA entrance channel (labeled as an asterisk). **D**) DHX29 model predicted by AlphaFold2 (**silver**), and model refined in the 43S cryo-EM density (**blue, purple, red**) showing the rearrangement of unstructured loops needed to place them in the density. **E)** The proposed model of the activity of the large insert upon ribosomal association. The different shades of pink represent heterogeneity in the position of the RecA2 domain in the left panel compared to the right.

While the resolution is low in this part of the complex, the eIF3g RRM appears to interact with DHX29’s CTR and to form part of DHX29’s mRNA channel entrance (**Figure 5A**), potentially facilitating the recruitment of mRNA to DHX29. The mRNA-binding activity of yeast eIF3g is mediated by the RRM (Hanachi et al., 1999), and eIF3g in human and C. elegans preferentially associates with mRNAs that have longer and more GC-enriched 5’UTRs than average (Lee et al., 2015; Blazie et al., 2021). These observations support a report that in yeast, mutations in the eIF3g RRM impaired translation of a mRNA containing a stable stem-loop in the 5’UTR by four-fold, consistent with a defect in scanning (Cuchalova et al., 2010). While it was long thought that DHX29 was only present in higher eukaryotes, a gene homologous to DHX29 and coding for a protein that interacts with the ribosome has recently been identified in yeast (Fromont-Racine, 2024). Taken together, these observations suggest that the eIF3g RRM and DHX29 may have synergistic roles in unwinding structured mRNAs. Our model will help generate more precise testable hypotheses to assess the importance and function of these interactions.

### MECHANISM OF AUTO-INHIBITION BY THE LARGE INSERT

The RecA2 domain of RHA-type helicases contains a large insert between the β1 strand and α1 helix (Dhote, 2012). Deletion of this domain does not affect DHX29’s function in promoting 48S complex formation and accurate start codon recognition but greatly increases DHX29’s basal NTPase activity when not bound to the ribosome or U_70_ RNA (Dhote 2012). The mechanism by which this large insert in the RecA2 domain inhibits the intrinsic NTPase activity of DHX29 remains elusive. Alphafold2 predicts the large insert forms a long β-hairpin that does not interact with any other domain of DHX29 **(Figure 1B, 1D, 3D)**. We observe an elongated density in the DHX29-bound 43S map at the entrance of the ribosome mRNA channel whose size and shape correspond well to that structure (**Figure 3B, 3C**). The rearrangement of unstructured loops that connect the large insert to the RecA2 domain allows us to fit the β-hairpin to this density with reasonable confidence, the length of these loops allowing the β-hairpin to reach that position (**Figure 3B, 3C, 3D**). The resolution in this part of the map is too low to detail specific interactions, but the large insert is in a position where it may form interactions with ribosomal proteins uS3, the NTT and CTT of uS5, and eS30, as well as with the RNA-recognition motif of eIF3g (**Figure 3B, 3C; S4**). Interestingly, this positions the β-hairpin of the large insert deeply into the mRNA channel, with the conserved K(A/G)G residues at its tip in position to interact either with the mRNA or with eS30 (**Figure S9**). This observation suggests that the β-hairpin may serve either as an mRNA sensor during scanning or to block the mRNA entrance channel and direct the mRNA towards DHX29’s RNA channel (**Figure 3B, 6A**).

Since we do not have a structure of DHX29 free in solution, we cannot infer directly how the large insert inhibits DHX29’s NTPase activity when not bound to the ribosome. Still, the AlphaFold2 model and our structure allow us to propose some hypotheses for a mechanism. Correct positioning of the RecA2 domain with respect to RecA1 appears key to the NTPase activity of DEAH helicases (Martin, 2013; Tauchert, 2017). The large insert and RecA2 domain being directly connected, their position and interactions are very likely to influence each other and, in turn, directly influence the NTPase activity.

One could hypothesize that the conformation of DHX29 predicted by Alphafold2, where the large insert does not interact with other domains of the protein, represents the conformation of the protein free in solution. In such a case, the autoinhibitory function of the domain would derive from the fact that, when DHX29 is free in solution, the RecA2 domain would be very dynamic and not stabilized in a productive conformation. We do not favor this hypothesis as it does not explain how mutation of the glutamine stretch relieves this autoinhibition and why deletion of the domain does not appear to affect DHX29 function (Dhote, 2012). An alternative hypothesis is that, when DHX29 is free in solution, the large insert is folded back onto the helicase core by interacting its highly conserved polyglutamine stretch with one of the positively charged surfaces of the helicase core (**Figure S9**). This position of the large insert would force the RecA2 domain into a conformation where catalysis cannot occur. The ribosome or U_70_ RNA would compete with this interaction, releasing the RecA2 from its unproductive conformation and stimulating NTPase activity.

We note that in several of the crystal structures of RHA family helicases, the structure of the large insert is not resolved (Schütz, 2010; Murakami, 2017; Chen, 2018), suggesting its flexibility in the absence of regulatory partners. Residues in the large insert of DHX9, DHX30, DHX36, and DHX57 do not share any sequence similarity with DHX29 (Dhote, 2012), and their structures show very different folds in Drosophila MLE (PDB: 5AOR), feline DHX9 (PDB: 8SZP), and in Alphafold2 predictions (**Figure S5**). This points to the large insert being specific to each helicase to accommodate their varied binding partners, while most likely still retaining similar regulatory functions by affecting the positioning of their RecA2 domain.

### THE RIBOSOME APPEARS TO STIMULATE DHX29 NTPASE ACTIVITY SIMILARLY TO A G PATCH PROTEIN

The ribosome stimulates DHX29’s NTPase activity by unknown mechanisms (Pisareva, 2008) and that stimulation is abrogated by mutations or deletions in both the NTR and the OB domain (Dhote, 2012; Sweeney, 2021). The map resolution in the region of the RecA domains (∼9-12 Å) only allowed us to refine the position of the RecA1 and RecA2 domains into the density of DHX29 bound to the 43S PIC as rigid bodies. To gain information on the state of the helicase core, we compared our model to the crystal structures of paralogs in different states. Our comparison suggests that the helicase core is in a conformation resembling that of Prp22 with ssRNA bound but without NTP (**Figure S6**) (Hamann, 2019), which presents a closed mRNA channel while the two RecA domains do not contact each other. Given that the reconstituted system used in the cryo-EM study contained the GTP analog GDPNP and no mRNA (Hashem, 2013), we would have expected the helicase domain to be in a state more similar to the structure of Prp43 with ATP analog ADP•BeF3^-^ and without mRNA (**Figure S6;** Tauchert, 2017), where the NTP binding pocket is closed, and the RNA channel is in an open state. This suggests that DHX29 is additionally regulated by other components in our reconstituted system, triggering the closed conformation of the RNA binding tunnel.

Adaptor proteins in the G patch family, named after a conserved short glycine-rich motif, regulate the activity and conformation of some members of the DEAH box RNA helicase family (Studer, 2020; Bohnsack, 2021). One such example is the protein NKRF, which regulates the function of DHX15, the mammalian analog of Prp43. The G patch motif of NKRF binds to DHX15, bridging its RecA2 and WH domains (Studer, 2020). By acting as braces, this interaction increases RNA affinity and promotes an open conformation of the ATP-binding pocket, enhancing both ATP turnover and processivity (Enders, 2022; Becker and Hub, 2023). Strikingly, h16 of the 40S subunit associates with DHX29 via an interface that overlaps significantly with the NKRF binding interfaces on DHX15 (**Figure 4B, 4C**). Our model shows that h16 interacts with the a.a.956-958 loop at the tip of a β hairpin called 5’-HP in the RecA2 domain, helix 1043-1056 of the WH domain, and the OB domain (**Figure 4A**). As anticipated, most residues at this interface are evolutionarily conserved across species encoding DHX29 (**Figure S7, S9**). In addition, the RecA2 domain of DHX29 is in a comparable position to that of DHX15 in complex with NKRF (**Figure 4A**). Based on these observations, we speculate that h16 may have a similar function to NKRF by stabilizing the RecA2 domain in a productive conformation (**Figure 4E)**. This would explain the upregulation of DHX29’s NTPase activity when it associates with the 40S subunit (Pisareva, 2008). The association with mRNA likely forms additional contacts that stabilize the RecA2 domain and further stimulate NTPase activity, as previously observed (Pisareva, 2008).

**Figure 4:**
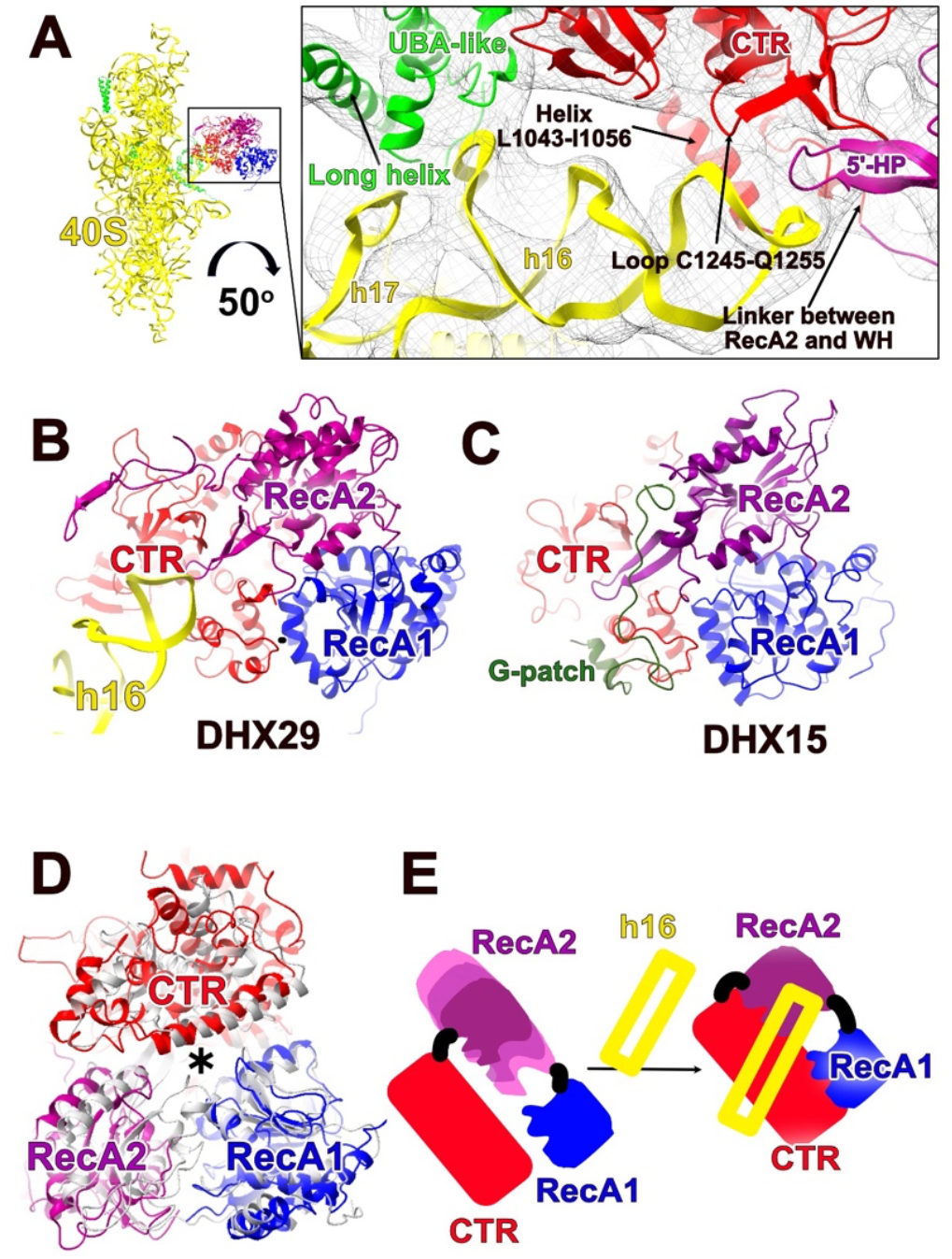
h16 may stimulate the NTPase activity of DHX29 similarly to a G-patch protein. **A**) h16 interaction with multiple domains of DHX29. **B**) A model of the regulation of h16 on the activity of DHX29 **C**) A model of the regulation of a G-patch protein on the activity of DHX15 **D)** Super-imposition of DHX15 bound with G-patch protein (**silver**, PDB: 6SH6) to the human DHX29 (**blue, purple, and red**) with the RecA1 domain (blue) as reference for alignment showing a similar conformation of the mRNA channel (labeled as an asteroid). (**E**) A schematic view of how the NTPase activity of h16 is enhanced by h16 of the 40S ribosome

### RESIDUES IMPORTANT FOR THE TRANSLOCASE ACTIVITY OF DEAH FAMILY HELICASES ARE CONSERVED

Whether DHX29 has intrinsic helicase activity or stimulates the helicase activity of the ribosome remains an open question. While DHX29 has not been shown to act as a processive helicase on its own (Pisareva, 2008), we cannot rule out the possibility that DHX29 itself unwinds the mRNA stem-loops in collaboration with the 40S subunit. To assess whether indeed DHX29 functions as a translocase, similarly to Prp22 and Prp43, we analyzed the position and the conservation of motifs critical for the function of these closely related DEAH helicases.

DExH-box helicases have been shown to translocate ssRNA through a channel formed between the helicase core and the CTR (Caruthers and McKay, 2002). We observe that the overall surface of the putative ssRNA channel of DHX29 is positively charged, the channel’s position and width resembling that of other DExH-box RNA helicases (**Figure 4D and S8**). It therefore appears likely to be a functional mRNA channel.

In the ATP-bound Prp22, a closed conformation of the helicase core has been observed, which allows motifs Ia, Ib, IV, and V as well as the hook-loop, the hook-turn, and the 5’-HP to interact with the ssRNA phosphate backbone in a sequence-independent manner (Hamann, 2019). Of the 19 residues that interact with the ssRNA backbone in Prp22, 9 are strictly conserved in DHX29 (**Figure 5**). The residues conserved in DHX29 include R1281 from the OB domain, R627 from motif Ia, R664, T679 from motif Ib, and R685, S806, T831 as well as N832 from motif V and K952, implying that DHX29 interacts with ssRNA via its phosphate backbone in a similar sequence-independent manner as Prp22.

**Figure 5:**
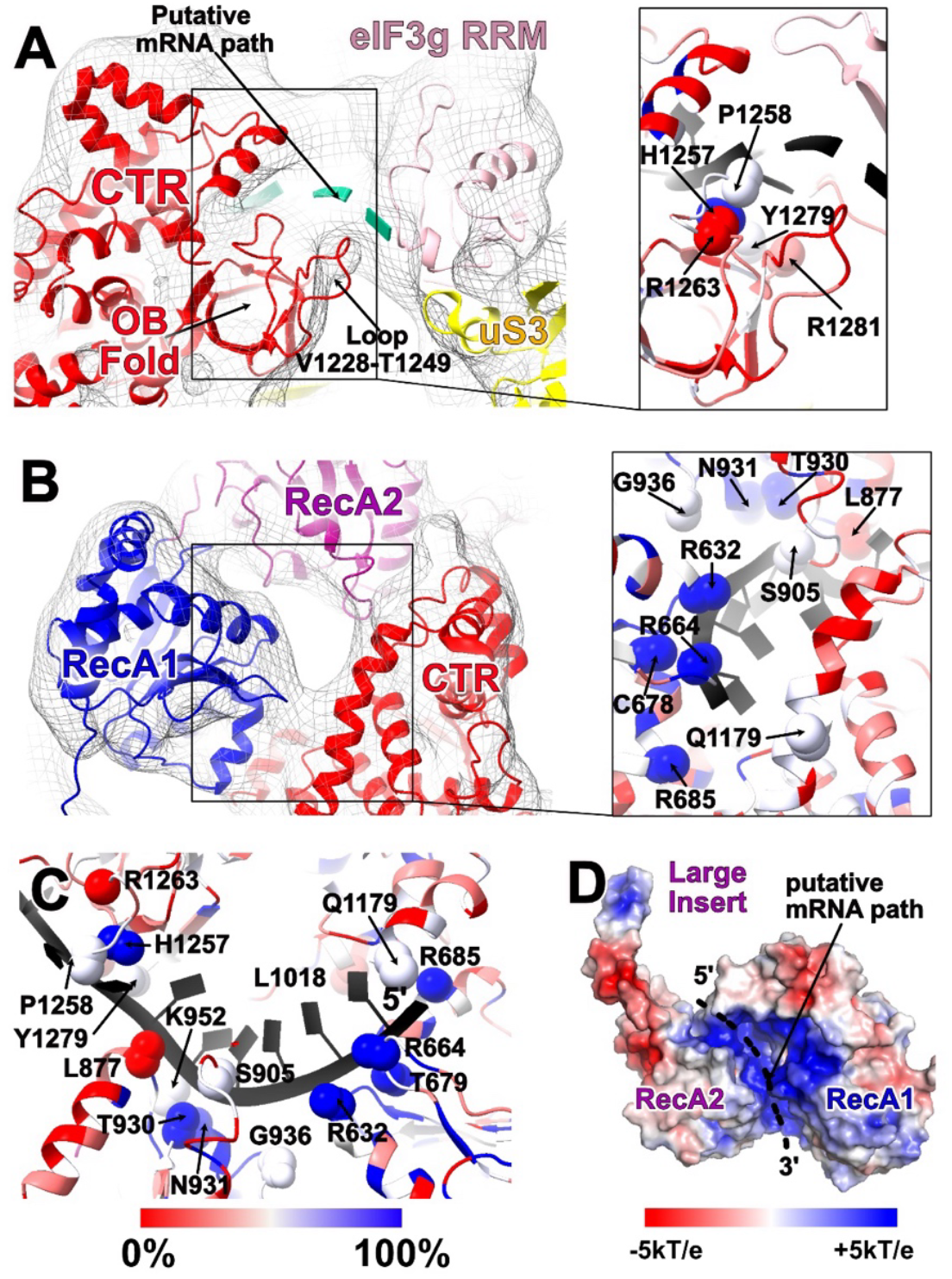
Sequence conservation of DHX29 RNA binding channel. **A**) Putative path of ssRNA based on the ssRNA-bound structure of prp22 (PDB: 6I3P), in proximity to eIF3g RNA recognition motif (RRM). The β-hairpin loop 1228-1249 of the OB domain and eIF3g RRM form part of the 5’ end of DHX29 putative mRNA channel.

As previously noted, the motifs I, II, and VI, which are critical for NTP binding and hydrolysis, are conserved in DHX29 (Dhote, 2012). We cannot readily assess the presence of GDPNP in the DHX29-bound 43S map due to the limited resolution, but the open conformation of the NTP binding pocket suggests the absence of GDPNP even though GDPNP was included in the reconstituted system (**Figure S6**). Motif III, which couples the NTPase activity to the helicase activity (Banroques, 2010), is strictly conserved among DExH-box helicases including DHX29, indicating that DHX29 may share a similar NTPase work cycle.

In addition to motif III, the hook-turn motif in the RecA1 domain is essential for RNA translocation in MLE (the Drosophila helicase maleless), Prp22, and Prp43 (Prabu, 2015; Tauchert, 2017; Hamann, 2019). The hook turn is conserved across all DExH-box helicases that bind RNA in a sequence-independent manner, but not in those that bind RNA in a sequence-dependent manner (Tauchert, 2017; Hamann, 2019). The fact that the first arginine residue in this motif is conserved in DHX29 (R664) and that the hook-turn of DHX29 is positioned adjacent to the putative mRNA path strongly suggest that the hook-turn has a similar function in DHX29 as in its paralogs (**Figure 5C**). If this assumption is correct, mutation of the hook-turn motif of DHX29 should abrogate RNA-dependent stimulation of NTPase activity similarly to Prp43 (Tanaka, 2006). In the RecA2 domain of SF2 DNA helicase Hel308, the 5’-HP motif is suggested to be involved in double-strand separation (Richards, 2008), but the function may not be conserved in DExH-box helicases given the different conformations of 5’-HP observed in Prp22 and Prp43 compared to Hel308 (He, 2010; Walbott, 2010; Tauchert, 2017; Hamann, 2019). The residues in 5’-HP in DHX29 that potentially interact with ssRNA are in a conformation more resembling Prp43 and Prp22 than Hel308 (**Figure 5C)**, suggesting that 5’-HP of DHX29 does not participate in unwinding activity either. Interestingly, the tip of the 5’-HP comes instead in direct head-on contact with h16 of the ribosome (**Figure 5A**). It is tempting to speculate that force applied by 5’-HP onto h16 during the helicase work cycle could induce conformational changes in the ribosome that participate in stem-loop unwinding.

On the 5’ side of the mRNA channel, DHX29 contains a conserved β-hairpin loop (V1228-T1249) potentially interacting with mRNA (**Figure 5A**), which is divergent from other DExH helicases. Overall, there is a higher level of conservation of residues between paralogs at the 3’end of the ssRNA channel (residues 80% conserved) compared to the 5’ side (residues 20% conserved). This may illustrate the divergent mode of anchoring to the substrate of DExH helicases in the 5’ region while residues implicated in the translocation mechanism appear conserved. Overall, our structural model suggests that DHX29’s mRNA channel is functional, interacting with mRNA in a sequence-independent manner. It shares key sequence and structural motifs with known 3’ to 5’ translocases such as Prp22 and Prp43, suggesting that DHX29 translocates ssRNA in a 3’ to 5’ direction. Still, this remains to be confirmed experimentally.

### DHX29 MAY ADOPT A SIMILAR MECHANISM AS ITS PARALOGS TO UNWIND STRUCTURED MRNAS

In the 43S PIC, DHX29 interacts with h16 of the ribosome, with its helicase RNA channel oriented towards the ribosomal mRNA channel (**Figure 6A**). As discussed above, given its structural similarity to DEAH-box helicases such as Prp22 and Prp43, DHX29 may employ a comparable 3’ to 5’ translocation mechanism. If so, this would present an intriguing issue: DHX29’s translocation direction would be opposite to the scanning direction.

**Figure 6:**
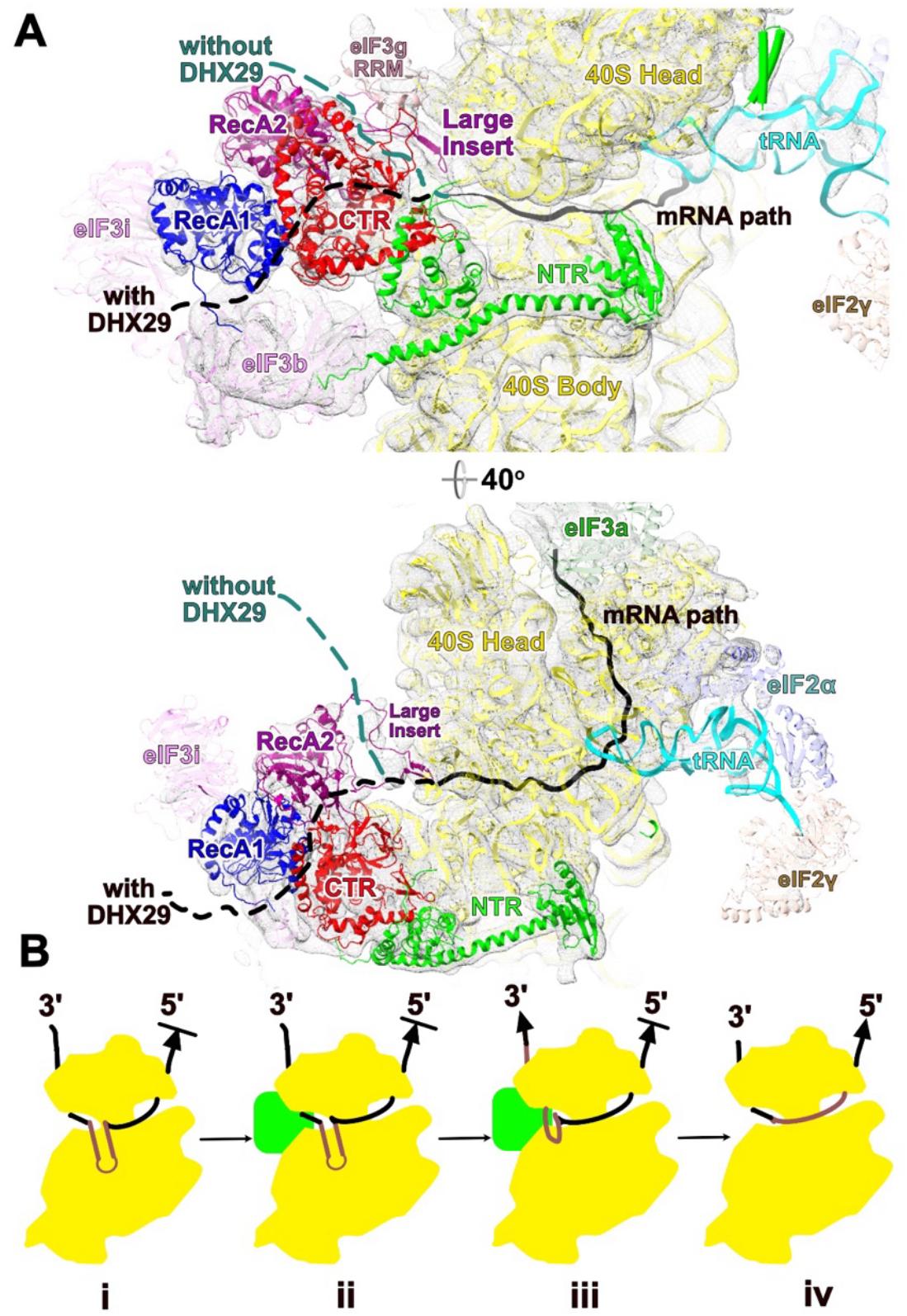
Model of stem-loop unwinding by DHX29. (**A**) A comparison of putative mRNA path during translation initiation upon DHX29 association to the 40S ribosome (**B**) The proposed model of DHX29’s unwinding activity during eukaryotic translation initiation: (i) Stem-loops bypasses the mRNA entry channel and are stuck in the ribosomal mRNA channel (**ii**) DHX29 recognizes the 3’-end of stem-loops (**iii**) Stem-loops are pulled and wrenched by DHX29 (**iv**) Stem-loops are further unwound by the 40S ribosome and re-enters the ribosomal mRNA channel; scanning resumes.

While DEAH-box RNA helicases appear to function as translocases, pulling RNA in a 3’ to 5’ direction, their unwinding activity often requires additional partners such as the spliceosome or the ribosome. It has been proposed that these helicases function in concert with other factors via a winching mechanism, requiring stable attachment to other structures to unwind RNA stem loops (Semlow, 2016). One possibility is that DHX29 uses a similar mechanism, pulling on the mRNA near the ribosomal entry channel in opposition to the scanning direction, thereby helping the ribosome unwind stable stem-loops as the mRNA is scanned from 5’ to 3’. Such a mechanism would allow the unwinding of stem loops stuck near the eIF2•GTP/Met-tRNA_i_^Met^ ternary complex or at the mRNA exit channel without requiring disassembly of the 43S PIC (Abaeva, 2011). Notably, in the DHX29 bound 43S PIC, the globular domains of eIF1 and eIF1A are not visible, and the mRNA channel adopts a closed conformation.

The dsRBD of DHX29 in our model partially overlaps with the expected position of eIF1A’s OB domain in the A site, suggesting a steric clash that may explain the absence of eIF1 and eIF1A in DHX29-bound complexes. This displacement of eIF1A by DHX29 likely contributes to mRNA channel closure in the DHX29-bound complex as eIF1A is needed to maintain an open mRNA channel (Passmore et al., 2007). In our proposed winching mechanism, this transient closure of the mRNA channel would be advantageous, as mRNA channel closure is unsuitable for scanning. In this state, the initiator tRNA may anchor the mRNA at the P-site, allowing the DHX29 helicase domain to pull on the mRNA stem-loop efficiently while preventing retrograde movement of the mRNA at the P-site. This would generate sufficient destabilization of the stem-loops for them to be unwound. Given that DHX29 is a non-processive helicase (Pisareva, 2008), it may release the mRNA after translocating a few nucleotides, allowing scanning to resume (**Figure 6B**).

Competition between eIF1A and DHX29 may play a crucial role in this process as in this model DHX29 would need to be displaced for eIF1A to bind, reopening the mRNA channel and enabling scanning to resume. It is important to note that only eIF1A’s globular domain appears to clash with DHX29. eIF1A may still remain attached to the 43S via its unstructured tails, allowing it to retain some of its functionalities and to efficiently compete with DHX29’s dsRBD. DHX29 appears to contain two dsRNA binding domains in its N-terminus, an HTH motif and a dsRBD domain, both positioned near the A-site. It is tempting to speculate that when an mRNA stem loop occupies the A-site, its recognition by these domains increases DHX29’s avidity for the 43S PIC, allowing it to outcompete eIF1A and consequently close the mRNA channel. Once the stem loop is unwound, it no longer stabilizes DHX29 in its site, reducing its avidity below that of eIF1A. This allows eIF1A to reoccupy the A-site, reopening the mRNA channel and enabling scanning to resume. This model would explain the drastic loss of function observed when the NTR of DHX29 is mutated or deleted (Sweeney, 2021).

However, an important limitation of this hypothesis is that no structure of a DHX29-bound preinitiation complex (PIC) containing mRNA has been resolved. We cannot rule out the possibility that eIF1A’s OB domain remains bound and the mRNA channel open under these conditions. Notably, Yi et al. (2022) observed eIF1A in an alternative conformation on an open 43S complex, a state that alleviates the steric clash with DHX29. Thus, direct competition between eIF1A’s OB domain and DHX29’s dsRBD for A-site binding remains to be confirmed.

Furthermore, additional mechanisms may contribute to stem-loop unwinding. DHX29’s NTPase cycle may generate force that propagates through the ribosome via the NTR, 5′HP, or its interactions with the 40S beak, leading to ribosome structural remodeling that stimulate its intrinsic helicase activity enough to scan through structured 5′UTRs, as suggested earlier by Pisareva et al. (2008). Similar mechanisms have been described for DEAH-box helicases, such as spliceosome ATPases, which induce conformational changes in large ribonucleoprotein (RNP) complexes to facilitate RNA remodeling (Tauchert et al., 2017; Semlow et al., 2016). It is also important to note that the backward pulling mechanism we propose here is not incompatible with some or all aberrant complexes requiring dissociation followed by new rounds of attachment and scanning as proposed by Abaeva et al. in 2011. Future mutagenesis and single-molecule fluorescence studies could help determine the precise nature of this mechanism.

## Conclusion

AlphaFold is a powerful tool for the prediction and validation of protein structures. The structure of human DHX29 was predicted by Alphafold2 with a high confidence level for most of the polypeptide chain, and is validated by its excellent agreement with the cryo-EM map of DHX29-bound 43S PIC. Refinements of the model based on the cryo-EM density allow us to infer new interactions between DHX29, the ribosome, and eIF3. Our model suggests that the ribosome helix 16 stimulates the NTPase activity of DHX29 similarly to a G-patch protein by stabilizing the position of several domains of DHX29. We also observe that the NTR of DHX29 bridges the body and head of the ribosome via unstructured linkers that could potentially transmit force between the ribosome and the helicase domain or constrain the mRNA path. The large insert of the RecA2 domain appears to slot deep into the mRNA channel and may serve as an mRNA sensor or direct it toward the helicase. Finally, most motifs found to be critical for the binding and translocation of mRNA in other DExH helicases appear conserved in DHX29, suggesting that it may be capable of mRNA translocation, potentially unwinding stem-loops via a winching mechanism. However, further experimental work is needed to determine whether DHX29 actively translocates along mRNA in a 3’ to 5’ direction or facilitates unwinding through ribosome-mediated structural changes. This model of DHX29 bound to 43S PIC offers an opportunity to generate new hypotheses and experiments to uncover the function and regulation of DHX29, a member of this intriguing family of DExH-box helicases.

## Acknowledgements

This research was supported by the National Institutes of Health (NIH) National Institute of General Medical Sciences (NIGMS) grants R35 GM122602 to T.V.P. and R01 GM097014 to C.H. and by startup funds provided by CUNY. The authors gratefully acknowledge Dr. Harsh Bansia and Dr. Daniel Keedy for their constructive input. We also thank Dr. Kevin Gardner for the thorough and critical review of the manuscript.

## Data Availability

The data underlying this article are available in the Protein Data Bank at www.rcsb.org, and can be accessed with PDB ID: 9CPA and at the Electron Microscopy Data Bank at www.ebi.ac.uk/emdb under EMDB ID: EMD-3057.

