## Supplementary Information for "The translation initiation factor DHX29 appears to pull on mRNA in a direction opposite to scanning"

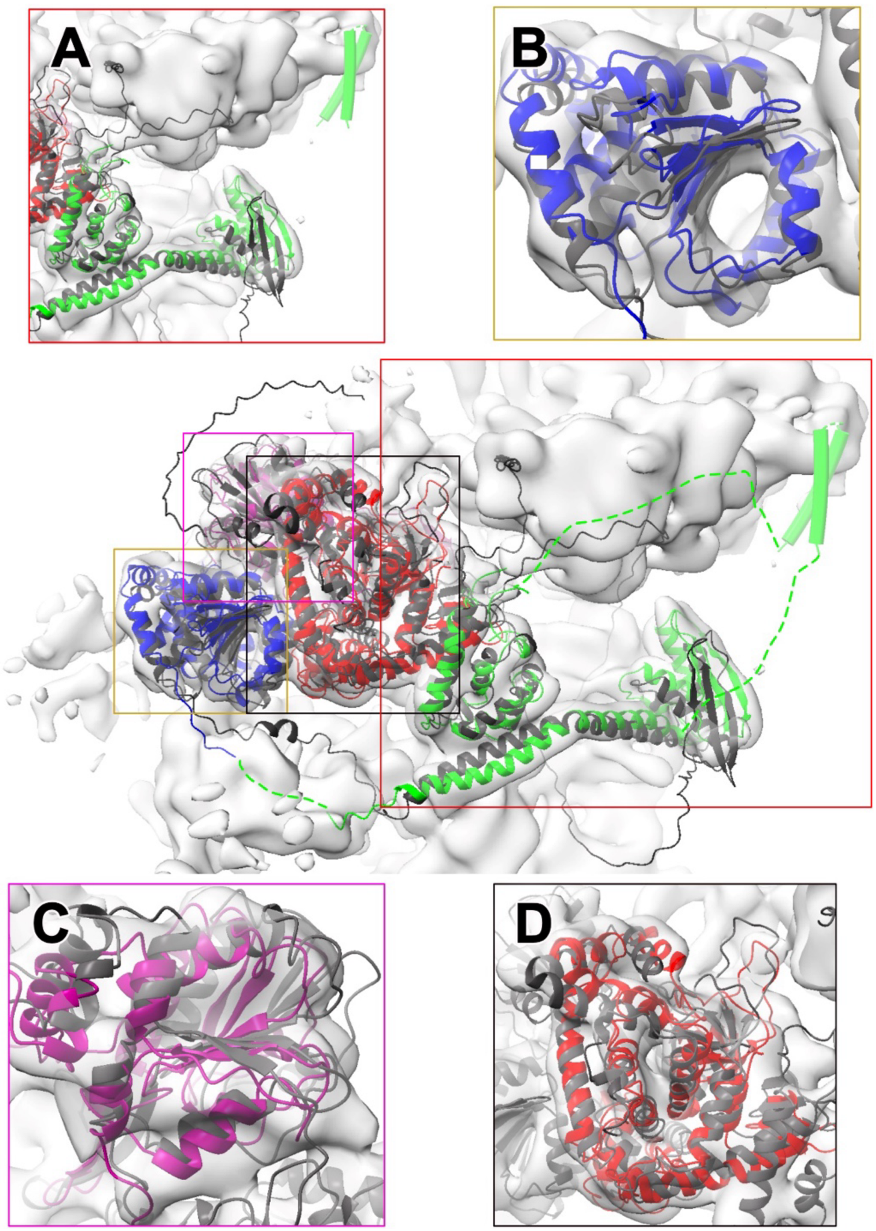


**Figure S1: Refinement of the AlphaFold model of human DHX29 into the 43S density**. Comparison between the DHX29 model predicted by Alphafold (light blue) and the final model refined in the DHX29 bound-43S density and colored by the domain (NTR: Green; RecA1: blue; RecA2: purple; CTR: red). Zoom in on the NTR (**A**), RecA1 **(B),** RecA2 (**C**), and CTR (**D**).

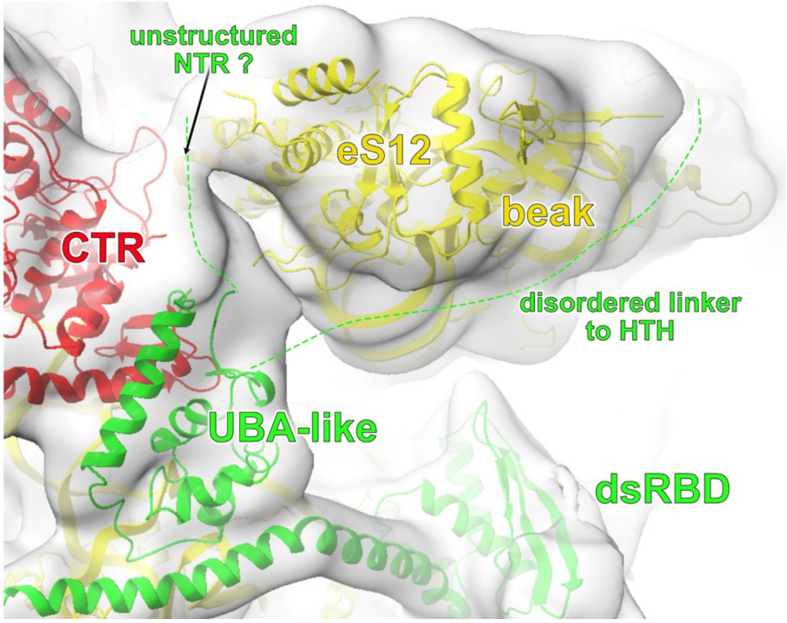


**Figure S2: Interaction between DHX29 and the ribosome beak**. Model of DHX29 bound to the ribosome zoomed on the beak region of the 40S ribosome and the NTR of DHX29 (green), overlaid with the 43S map filtered at 12Å resolution.

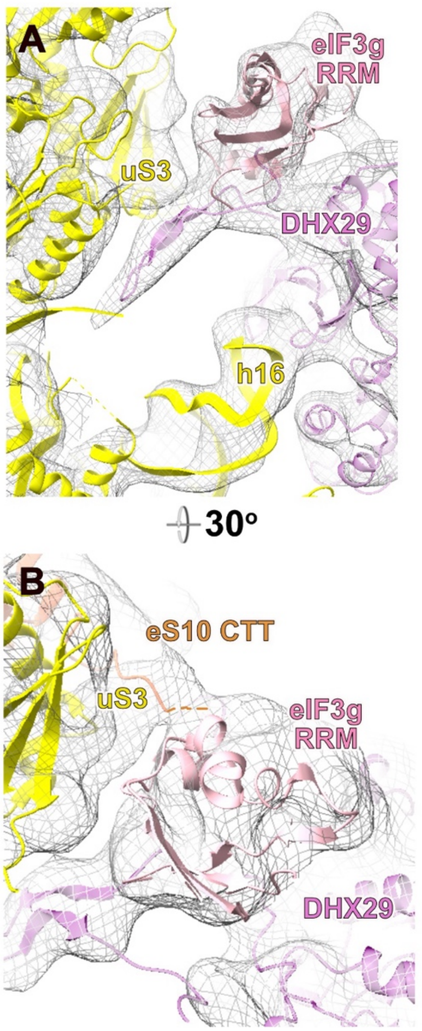


**Figure S3: Comparison of eIF3g RRM interaction with the 40S ribosome between DHX29-bound 43S and 48S without DHX29**. **(A)(B)** The interaction between eIF3g RRM and DHX29-bound mammalian 43S complex. The putative model for extension in eS10 CTT is depicted as a dotted line. **(C)(D)** The interaction between eIF3g RRM and human 48S complex with DHX29 association.

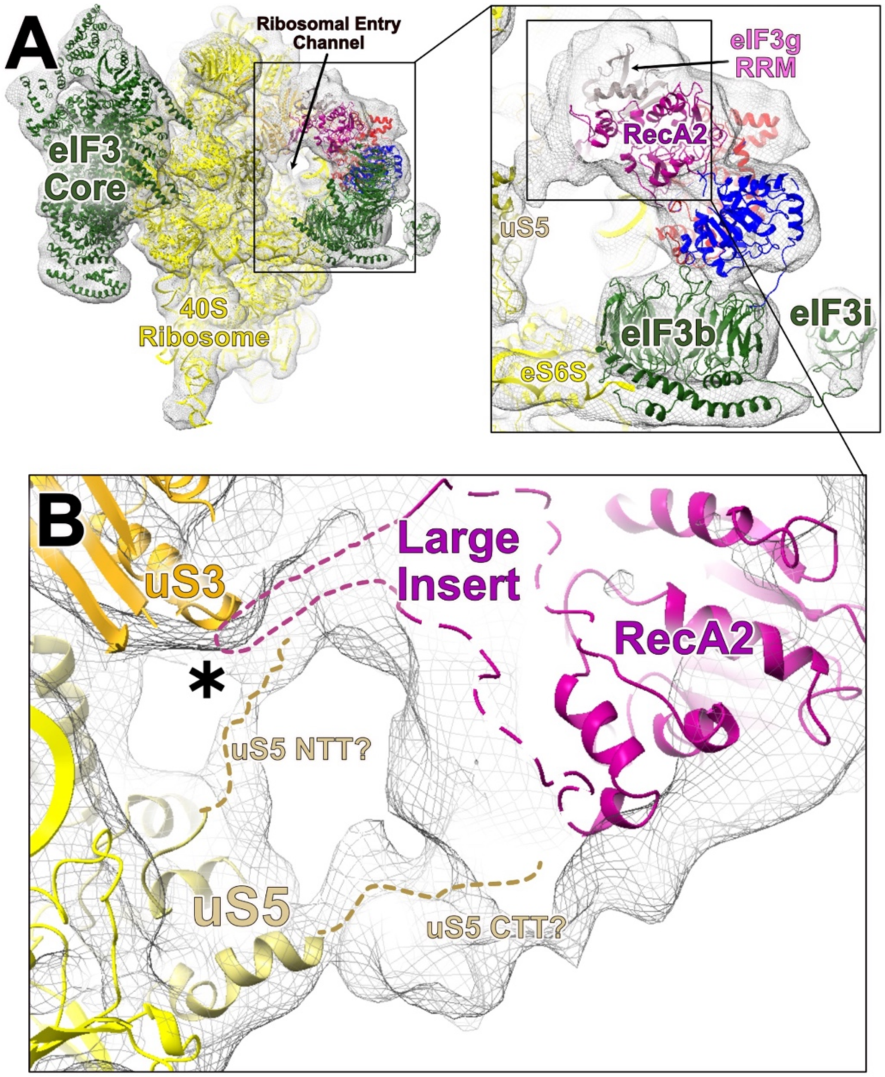


**Figure S4: Potential interaction of uS5 with the large insert of DHX29**. (**A**) An overview of the interaction network around the mRNA entry channel of DHX29-bound 43S complex (**B**) Low-resolution densities connecting uS5 and the large insert are observed. The dash lines represent the putative NTT and CTT of uS5 as well as the flexible linker in the large insert.

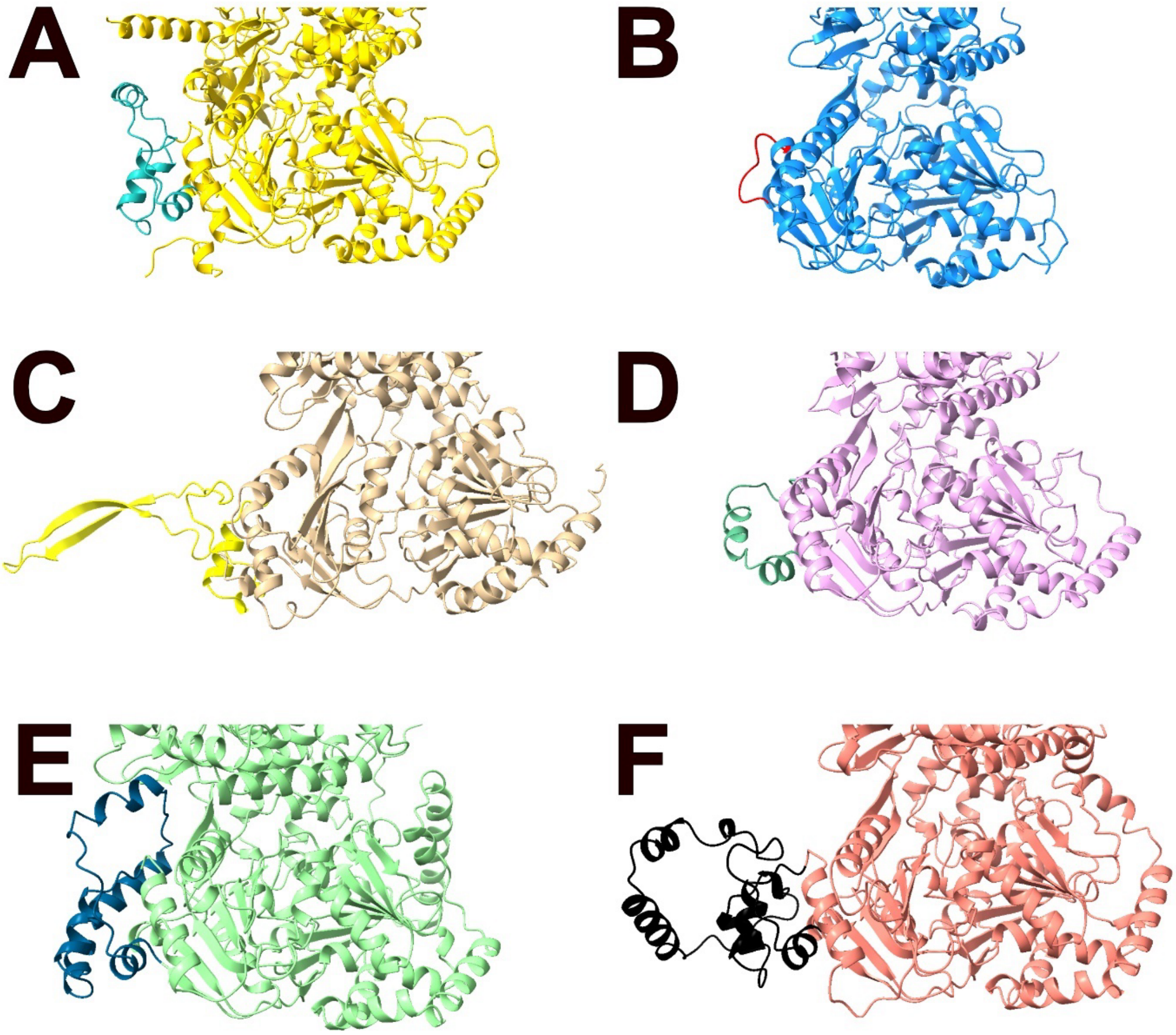


**Figure S5: Alphafold2-predicted structures of the large insert in different RHA helicases**. Only the rigid helicase core and the large insert are shown, and the large insert is separately colored. **A**) human DHX9. **B**) human DHX15. **C**) human DHX29. **D**) human DHX36. **E**) human DHX56. **F**) human DHX57.

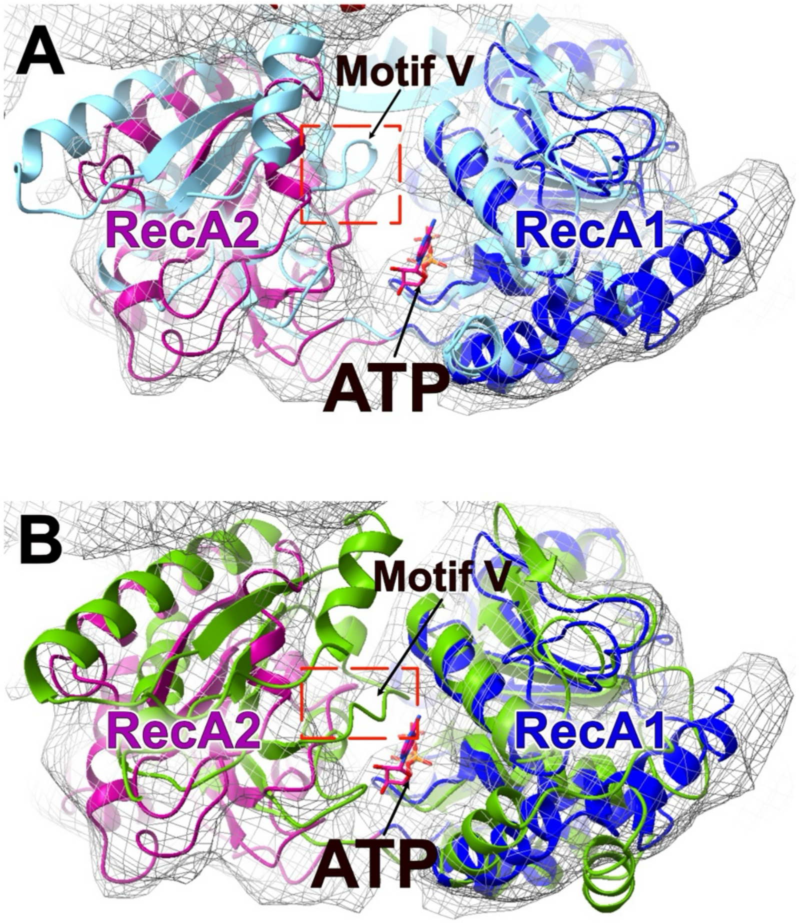


**Figure S6: Structural comparison of the NTP binding pocket between DHX29 and its paralogs**. **A**) Comparison of the NTP binding pocket between DHX29 (green and purple) and Prp22 bound with ssRNA and without NTP (light blue; PDB: 6I3P), when aligned to each other with RecA1 domain as reference. The ATP position is derived from the structure of Prp43 bound with ATP analog ADP•BeF3- (PDB: 5LTK). Motif V, which controls the opening and closing of the NTP binding pocket, is highlighted with a red box. **B**) Comparison of the NTP binding pocket between DHX29 (green and purple) and Prp43 bound with ATP analog ADP•BeF3- and without NTP (green; PDB: 5LTK).

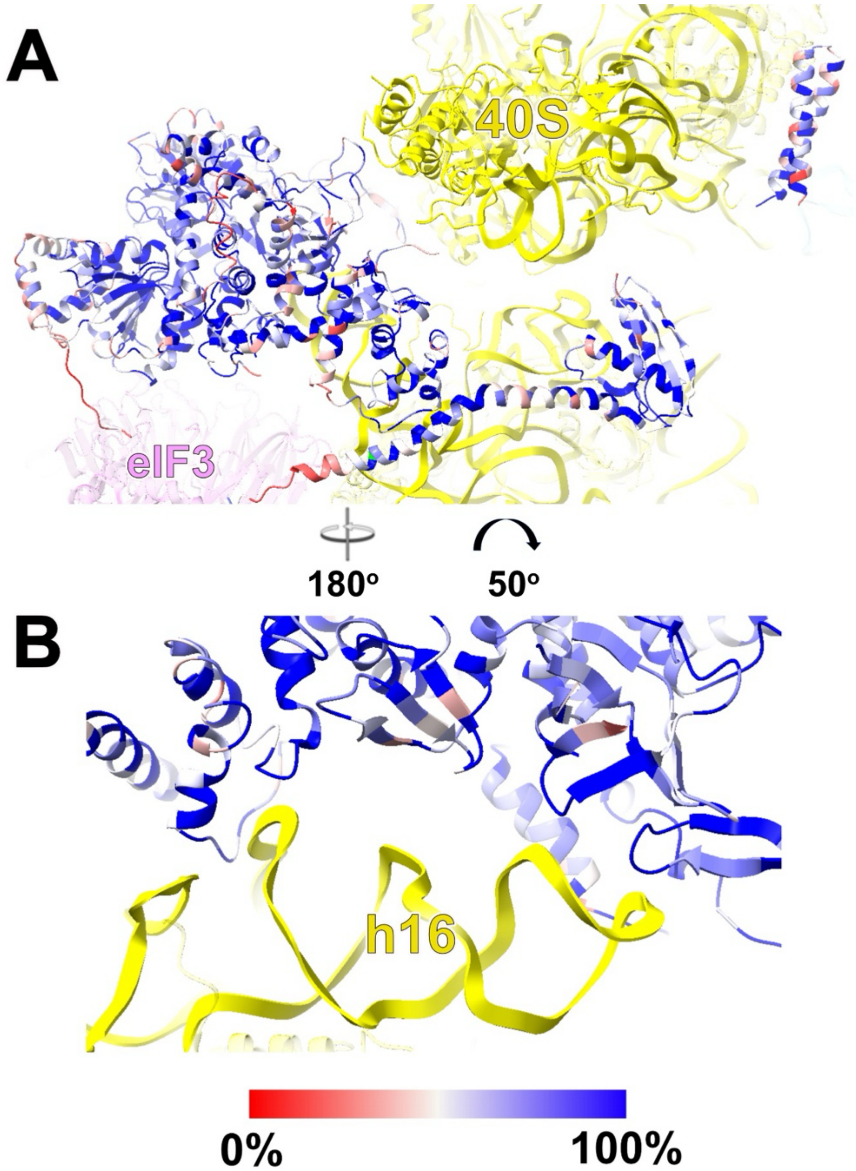


**Figure S7: Sequence conservation of DHX29**. Sequence conservation was measured across 21 organisms from yeast, Chordata, Mollusca, and Echinodermata. **A**) Sequence conservation of all DHX29 residues. **B)** Sequence conservation of residues interacting with h16.

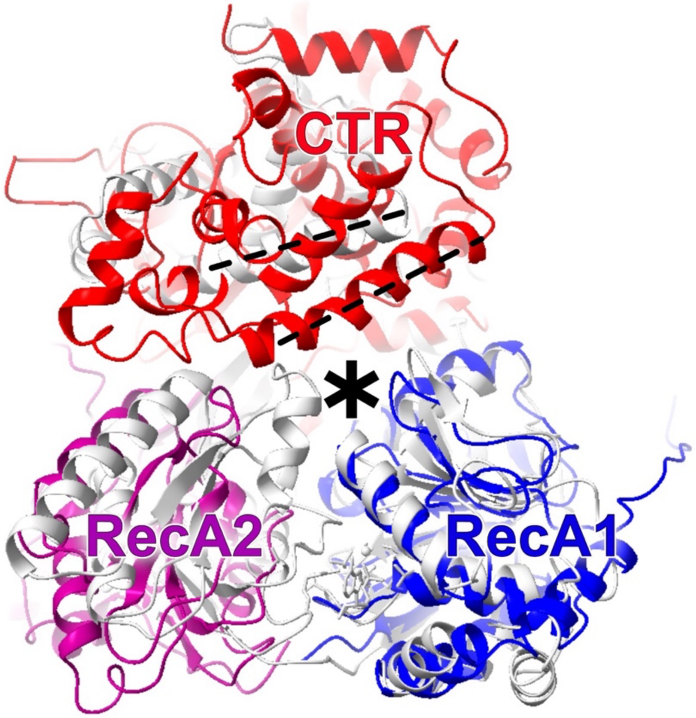


**Figure S8: Structural comparison between DHX29 and Prp43 bound with ATP analog ADP•BeF3- and without mRNA.** DHX29 (blue, purple, red) and Prp43 (dark grey, PDB: 5LTK) super-imposed onto each other with the RecA1 domain (blue) as alignment reference. The mRNA channel is marked with an asterisk.

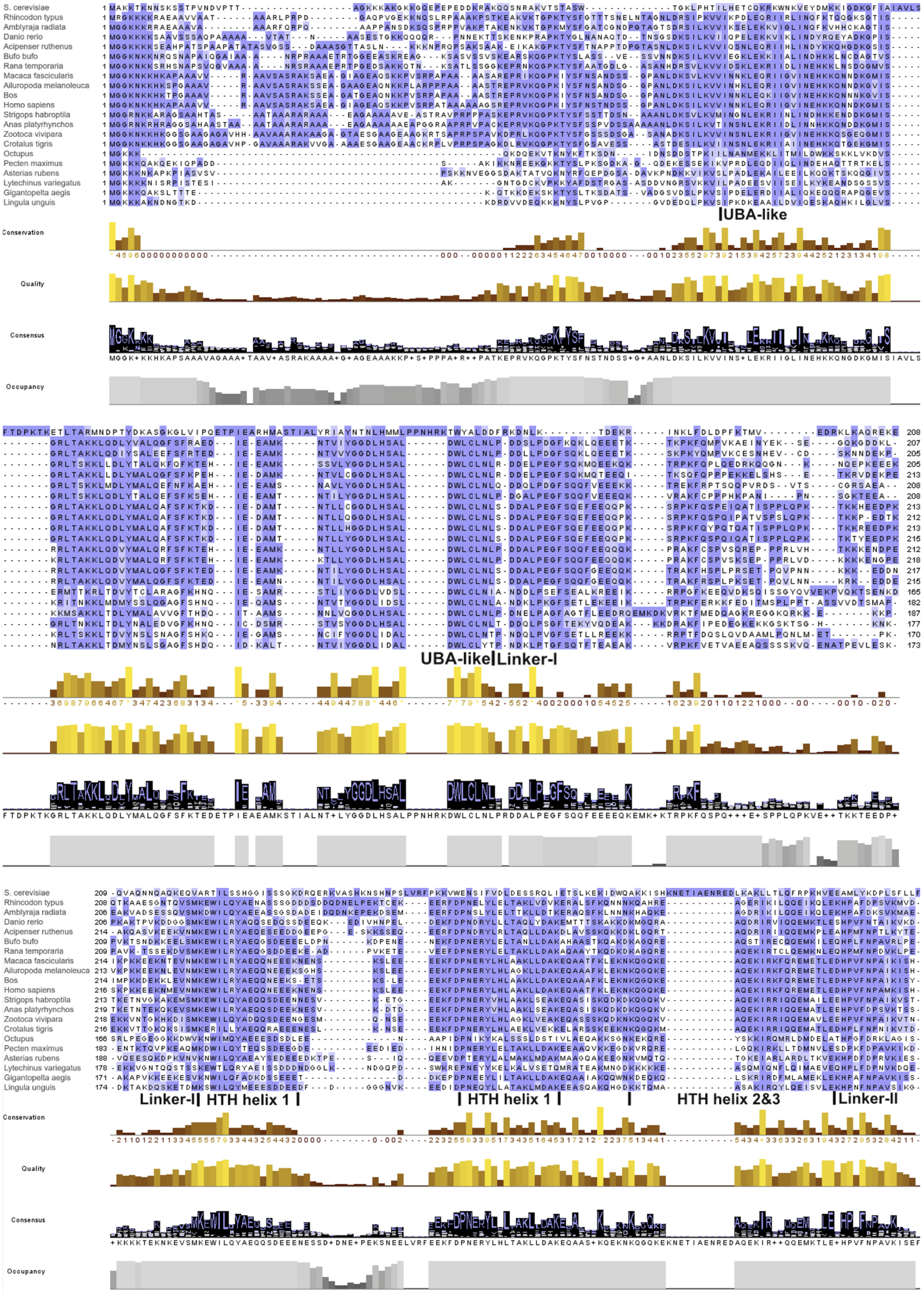
S. cerevisiae
Rhincodon typus
Amblyraja radiata
Danio rerio
Acipenser ruthenus
Bufo bufo
Rana temporaria
Macaca fascicularis
Ailuropoda melanoleuca
Bos
Homo sapiens
Strigops habroptila
Anas platyrhynchos
Zootoca vivipara
Crotalus tigris
Octupus
Pecten maximus
Asterias rubens
Lytechinus variegatus
Gigantopelta aegis
Lingula unguis
UBA-like
Conservation
Quality
Consensus
Occupancy
UBA-like
Linker-I
Linker-I
HTH helix 1
HTH helix 1
HTH helix 2&3
Linker-II

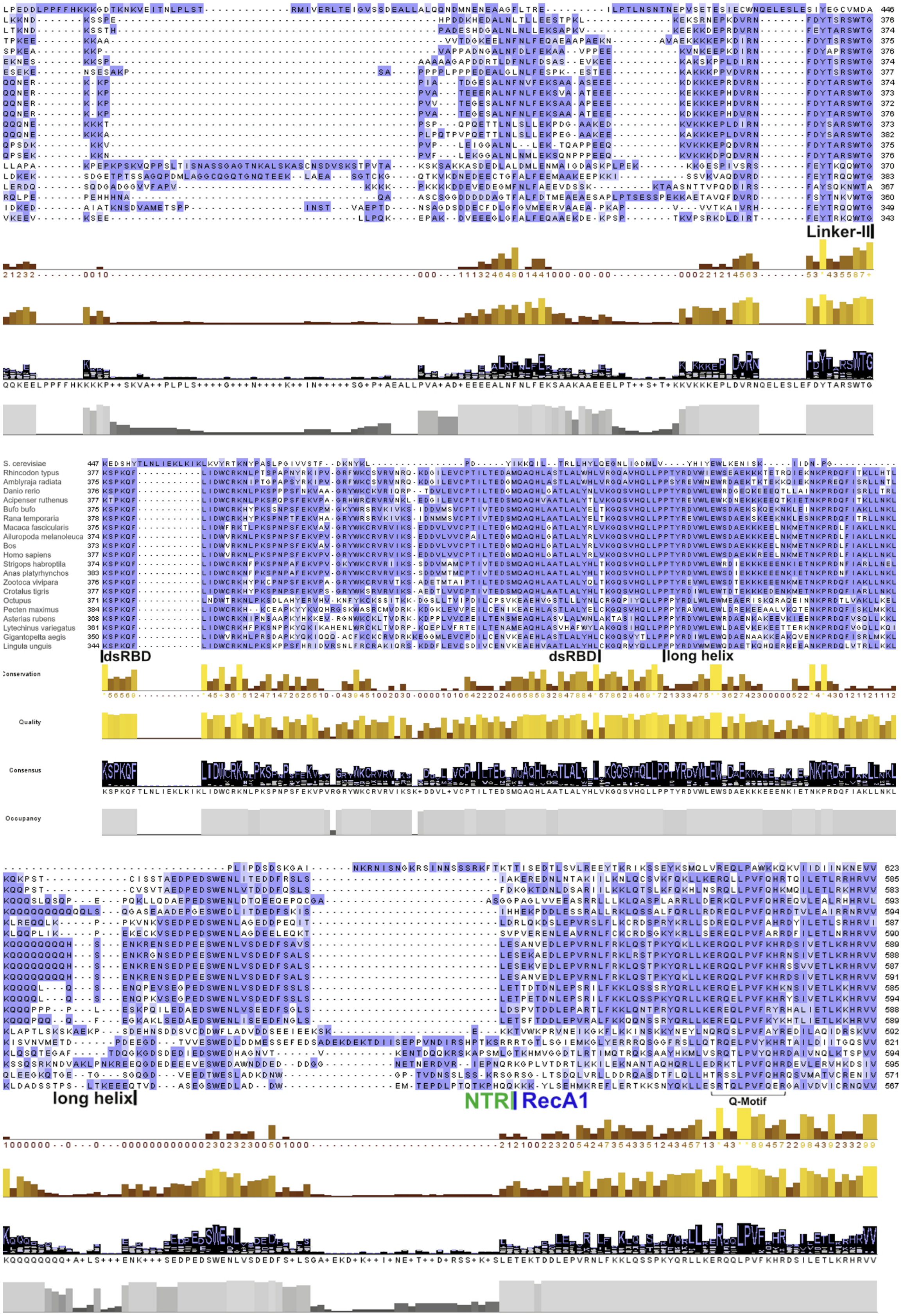
Linker-II
S. cerevisiae
Rhincodon typus
Amblyraja radiata
Danio rerio
Acipenser ruthenus
Bufo bufo
Rana temporaria
Macaca fascicularis
Ailuropoda melanoleuca
Bos
Homo sapiens
Strigops habroptila
Anas platyrhynchos
Zootoca vivipara
Crotalus tigris
Octupus
Pecten maximus
Asterias rubens
Lytechinus variegatus
Gigantopelta aegis
Lingula unguis
dsRBD
dsRBD
long helix
Conservation
Quality
Consensus
Occupancy
long helix
NTR
RecA1
Q-Motif

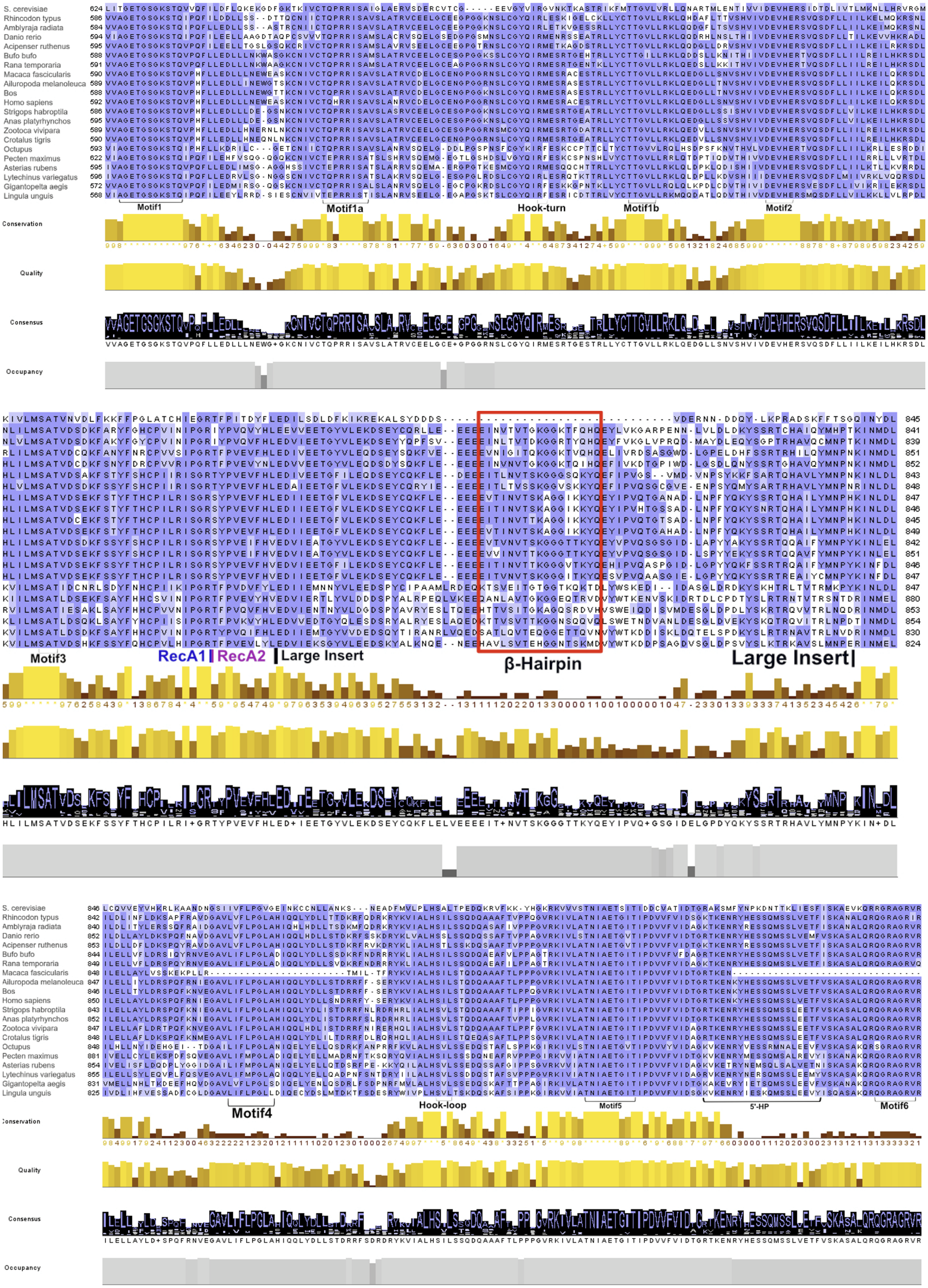
S. cerevisiae
Rhincodon typus
Amblyraja radiata
Danio rerio
Acipenser ruthenus
Bufo bufo
Rana temporaria
Macaca fascicularis
Ailuropoda melanoleuca
Bos
Homo sapiens
Strigops habroptila
Anas platyrhynchos
Zootoca vivipara
Crotalus tigris
Octupus
Pecten maximus
Asterias rubens
Lytechinus variegatus
Gigantopelta aegis
Lingula unguis
Motif1
Motif1a
Hook-turn
Motif1b
Motif2
Conservation
Quality
Consensus
Occupancy
Motif3
RecA1
RecA2
Large Insert
β-Hairpin
Large Insert
Motif4
Hook-loop
Motif5
5'-HP
Motif6

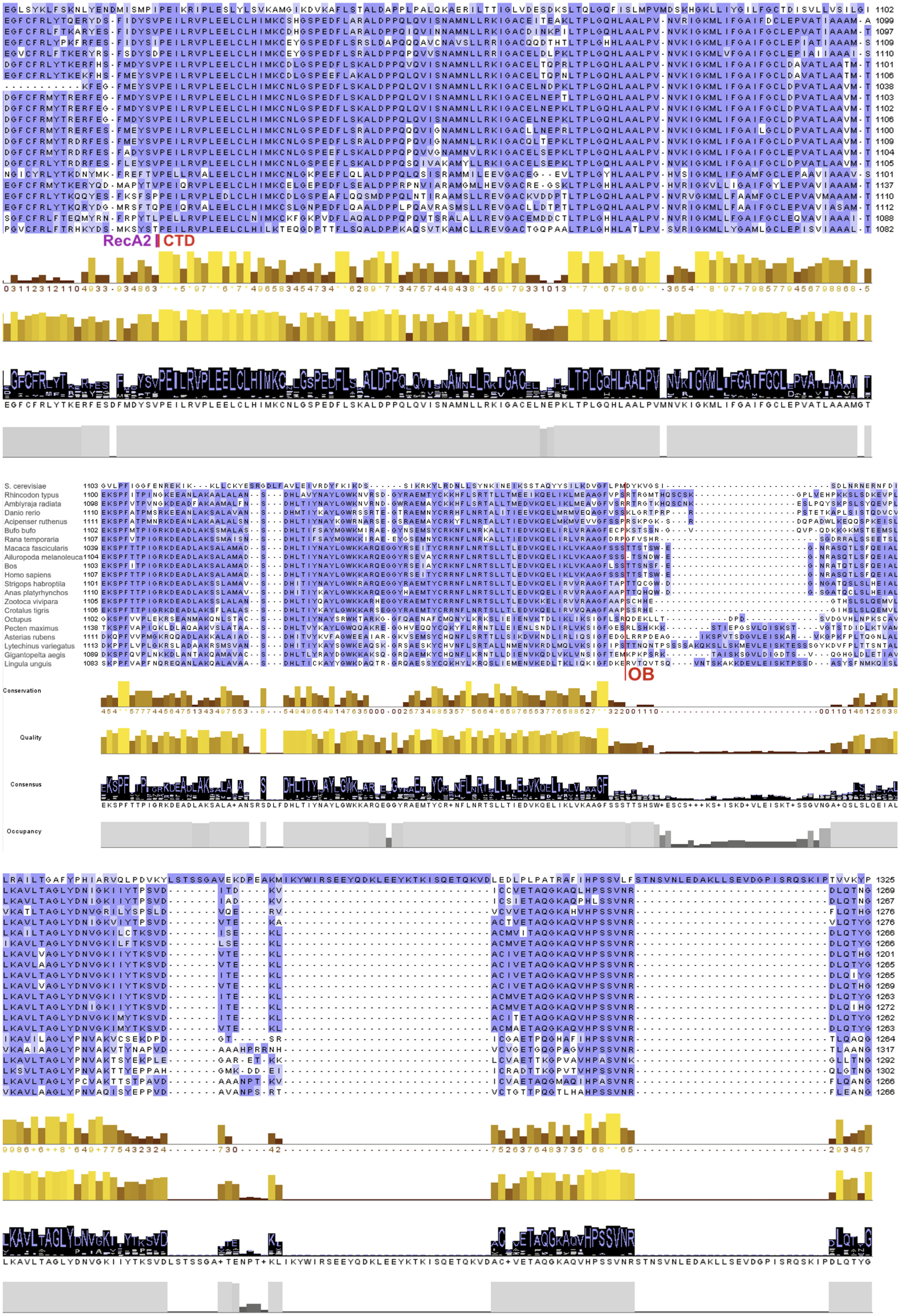
RecA2
CTD
S. cerevisiae
Rhincodon typus
Amblyraja radiata
Danio rerio
Acipenser ruthenus
Bufo bufo
Rana temporaria
Macaca fascicularis
Ailuropoda melanoleuca
Bos
Homo sapiens
Strigops habroptila
Anas platyrhynchos
Zootoca vivipara
Crotalus tigris
Octupus
Pecten maximus
Asterias rubens
Lytechinus variegatus
Gigantopelta aegis
Lingula unguis
OB
Conservation
Quality
Consensus
Occupancy

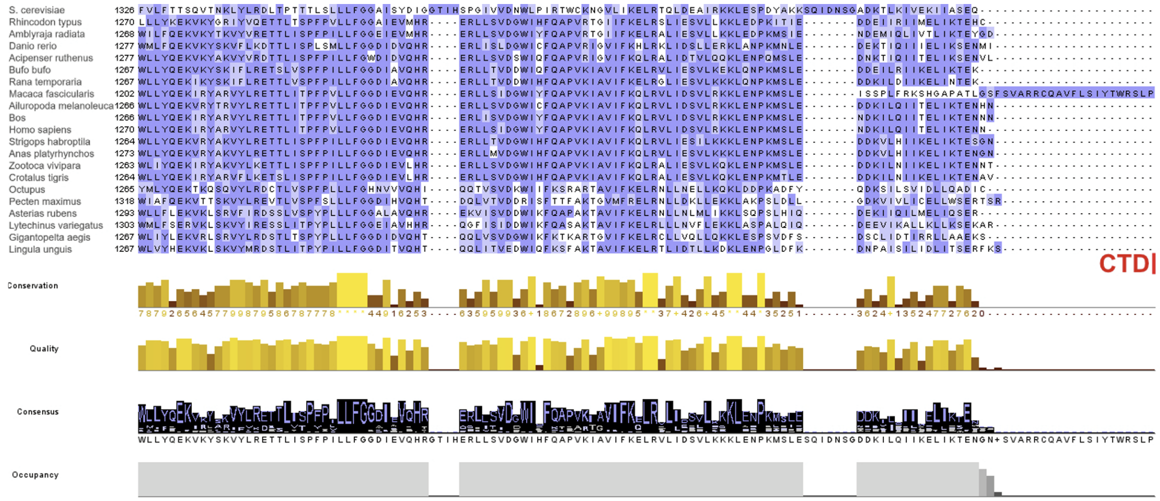


**Figure S9: Sequence alignment of DHX29 from 21 species**. Species are *S. cerevisiae* (Taxon: 559292), *Rhincodon typus* (Taxon: 259920), *Amblyraja radiata* (Taxon: 386614), *Danio rerio* (Taxon: 7955), *Acipenser ruthenus* (Taxon: 7906), *Bufo bufo* (Taxon: 8384), *Rana temporaria* (Taxon: 212803), *Macaca fascicularis* (Taxon: 3016450), *Ailuropoda melanoleuca* (Taxon: 9646), *Bos* (Taxon: 9903), *Homo sapiens* (Taxon: 9606), *Strigops habroptila* (Taxon: 2489341), *Anas platyrhynchos* (Taxon: 8840), *Zootoca vivipara* (Taxon: 2563720), *Crotalus tigris* (Taxon: 88082), *Octopus* (Taxon: 6610), *Pecten maximus* (Taxon: 6579), *Asterias rubens* (Taxon: 7604), *Lytechinus variegatus* (Taxon: 1316090), *Gigantopelta aegis* (Taxon: 1735272), and *Lingula unguis* (Taxon ID: 7574). Residues that are more than 90% conserved are colored deep blue; residues that are 65% to 90% conserved are colored cornflower blue; residues that are 50% to 65% conserved are labeled in light blue. Key motifs are label
